# Cyclin A and Cks1 promote kinase consensus switching to non-proline directed CDK1 phosphorylation

**DOI:** 10.1101/2022.05.24.493195

**Authors:** Aymen al-Rawi, Svitlana Korolchuk, Jane Endicott, Tony Ly

## Abstract

Ordered protein phosphorylation by CDKs is a key mechanism for regulating the cell cycle. How temporal order is enforced in mammalian cells remains unclear. Using a fixed cell kinase assay and phosphoproteomics, we show how CDK1 activity and non-catalytic CDK1 subunits contribute to the choice of substrate and site of phosphorylation. Increases in CDK1 activity alters substrate choice, with intermediate and low sensitivity CDK1 substrates enriched in DNA replication and mitotic functions, respectively. This activity dependence was shared between Cyclin A- and Cyclin B-CDK1. Cks1 has a proteome-wide role as an enhancer of multisite CDK1 phosphorylation. Contrary to the model of CDK1 as an exclusively proline-directed kinase, we show that Cyclin A and Cks1 promote non-proline directed phosphorylation, preferably on sites with a +3 lysine residue. Indeed, 70% of cell cycle regulated phosphorylations, where the kinase carrying out this modification has not been identified, are non-proline directed CDK1 sites.

## Introduction

Ordered phosphorylation by cyclin-dependent kinases (CDKs) controls the timing and progression of the cell division cycle. Temporal regulation of CDK phosphorylation is critical, ensuring that DNA replication origin licensing, DNA replication, and chromosome segregation occur in sequential order (Gavet and Pines, 2010; Morgan, 1997; Swaffer et al., 2016). CDK1 is essential to embryonic cell division and supports cell division in the absence of interphase CDKs (CDK2/4/6)(Santamaría et al., 2007). In *S. pombe*, a single cyclin-CDK fusion can drive a relatively unperturbed cell cycle in optimal growth conditions (Coudreuse and Nurse, 2010). How temporal ordering of CDK1 phosphorylation is controlled in mammalian cells remains a major open question.

Complex formation with a cognate cyclin is requisite to CDK activation. Mammalian genomes encode four major cell cycle CDKs (CDK1/2/4/6) and ten cyclins (D1/D2/D3/E1/E2/A1/A2/B1/B2/B3). Mouse knockout studies have elegantly shown the differential requirements of cyclins and CDKs in embryogenesis and development (Santamaría et al., 2007; Satyanarayana and Kaldis, 2009) . However, detailed biochemical understanding of how CDK phosphorylation is controlled in mammalian cells is hampered by redundancy in the cyclin-CDK family and kinase-phosphatase feedback loops (e.g., Wee1/Cdc25, Greatwall/PP2A-B55) that can be acutely sensitive to changes in CDK activity in cells (Hégarat et al., 2020; Lau et al., 2021; Mitra and Enders, 2004; Russell and Nurse, 1986, 1987).

Cks is a third subunit found in active cyclin-CDK complexes that is conserved from yeast (Suc1 in *S. pombe*) to human. Cks1 acts as a phospho-adaptor protein with selectivity for phosphothreonine (McGrath et al., 2013). Mammalian genomes encode two Cks proteins (CKS1B and CKS2), and deletion of both leads to embryonic lethality (Martinsson-Ahlzén et al., 2008). Cks1 (encoded by CKS1B), but not Cks2, promotes the ubiquitination and degradation of the CDK inhibitor protein p27^KIP^ and promotes cell cycle entry (Ganoth et al., 2001; Sitry et al., 2002). In *S. cerevisiae*, Cks1 facilitates multisite phosphorylation of the CDK substrate protein, Sic1 (Köivomägi et al., 2011). Docking of Cks1 onto priming sites is thought to promote the phosphorylation of low affinity sites by *cdc28* (CDK1 in human) (Kõivomägi et al., 2013). It is unclear how many CDK substrates require Cks1 for high occupancy phosphorylation.

In *S. pombe*, a quantitative model of CDK1 substrate choice was proposed whereby substrates with functions in DNA replication have lower thresholds for CDK1 phosphorylation than substrates with functions in mitosis (Fisher and Nurse, 1996; Stern and Nurse, 1996). Thus, temporal ordering is achieved by a progressive increase in CDK1 activity. In *S. cerevisiae*, qualitative differences, for example in the Cyclin subunit, confers substrate specificity to CDK1 (Loog and Morgan, 2005; Örd and Loog, 2019; Örd et al., 2019). Cyclins E, A, and D have a hydrophobic patch ∼35 Å from the catalytic site that has been shown to be important in substrate recognition by binding to a Cy motif (RXL, where X is any amino acid) in disordered regions (Adams et al., 1996; Lowe et al., 2002; Schulman et al., 1998; Takeda et al., 2001). Mutation of the hydrophobic patch significantly reduces phosphorylation occupancy for a subset of substrates. The hydrophobic patch is not completely degenerate between cyclins because identical mutations have differential effects on substrate phosphorylation dependent on the Cyclin subunit (Loog and Morgan, 2005). Additionally, cyclins are targeted to distinct subcellular compartments (Jackman et al., 2002; Moore et al., 2002; Toyoshima et al., 1998). This targeting is encoded in nuclear localization and export sequences that differ between cyclins. Consistent with these results, cyclins have overlapping and complementary interaction partners in cells (Pagliuca et al., 2011). Cyclins also have non-redundant functions. Cyclin A2 is essential for mitotic entry and prevents hyperstable kinetochore-microtubule attachments in early mitosis (Gong et al., 2007; Hégarat et al., 2020; Kabeche and Compton, 2013). In contrast, cells depleted of Cyclin B proteins progress into mitosis, but fail to complete proper chromosome segregation (Hégarat et al., 2020).

CDK1 phosphorylation of individual sites on a substrate protein is ordered to produce ultrasensitive switches in protein function (Trunnell et al., 2011). However, little is known about how this order is regulated. Phosphorylation site choice within a substrate is determined by molecular interfaces proximal to the catalytic site. A structure for Cyclin A-CDK2 in complex with a peptide substrate shows that a substrate binding cleft formed on one side by the activation loop imposes a preference for a proline in the +1 position and a basic residue in the +3 position (Brown et al., 2015; Brown et al., 1999a). These structural studies are consistent with data from peptide arrays and biochemical assays showing a consensus sequence of [ST]PX[KR] for *cdc28*, which is a serine or threonine, followed by a proline and a basic residue (lysine or arginine) in the +3 position (Mok et al.). Compared to CDK2, the activation loop of CDK1 is more flexible, possibly allowing for a relaxed consensus requirement. Consistent with this idea, CDK1 phosphorylates synthetic peptides with non-optimal consensus sequences that lack a proline in the +1 position, but only if the peptide contains a RXL motif (Brown et al., 2015). By contrast, under the same reaction conditions, CDK2 is unable to phosphorylate these ‘non-canonical’ CDK sites. Non-canonical CDK1 sites have been described for individual protein substrates and for substrates in mESCs (Kõivomägi et al., 2013; Michowski et al., 2020; Suzuki et al., 2015). However, it is unknown what proportion of the CDK substrate phosphorylations are on non-canonical sites, and if and how non-canonical phosphorylations are regulated.

Recent advances in mass spectrometry (MS)-based proteomics have enabled quantitative and comprehensive analysis of cellular protein phosphorylation (Dephoure et al., 2008; Herr et al., 2020; Olsen et al., 2010). These approaches have produced an extensive catalogue of phosphorylation sites in human cells that are cell cycle regulated, recently with high temporal resolution (Ly et al., 2017) . Many cell cycle regulated phosphorylation sites do not match any known kinase consensus sequence. As an example, 28% of the HeLa mitotic phosphoproteome do not match a cell cycle kinase consensus motif (CDK: [ST]P, Plk1: [DE]X[ST][Φ]X[DE], Aurora: [KNR]RX[ST][Φ], where Φ is a hydrophobic residue) (Dephoure et al., 2008). These sites with no predicted upstream kinase constitute a dark fraction of the cell cycle regulated phosphoproteome. Interestingly however, the sites are enriched in a motif consisting of a S/T followed by a K in the +3 position, leading to speculation that there are unknown kinases that drive a significant fraction of cell cycle regulated phosphorylation (Dephoure et al., 2008).

In this study, we developed an *in vitro* approach to investigate how quantitative and qualitative characteristics of the CDK1 complex contribute to substrate choice and phosphorylation. Fixed and permeabilized cells are subjected to kinase reactions with recombinant CDK1 in complex either with Cyclin B, Cyclin A, or Cyclin B-Cks1. We show that both Cyclin A2 and Cks1 promote the phosphorylation of non-proline sites by CDK1 *in vitro*. Cyclin A2-CDK1 non-proline directed sites are enriched in a lysine in the +3 position (+3K), in a motif defined by [ST][^P]XK, where [^P] indicates any residue except for proline. Sites detected *in vitro* are cell cycle regulated *in vivo*, including non-proline directed sites and sites containing the +3K. By combining sequential enzymatic reactions on fixed cells, we demonstrate that the majority of Cks1-promoted sites are primed by CDK1 itself. Based on our data, we propose a model whereby the cyclin subunit determines substrate specificity, whereas the role of Cks1 is an enhancer of cyclin-CDK1 activity by promoting multisite phosphorylation of low affinity sites.

## Results

### A fixed cell kinase assay to investigate substrate phosphorylation proteome-wide in vitro

To investigate the role of the Cks1 and the cyclin subunits in CDK1 substrate phosphorylation, we designed an *in vitro* kinase assay in which formaldehyde fixed and methanol permeabilized TK6 cells are incubated with purified CDK1 in complex with either Cyclin B1 (BC), Cyclin B1 and Cks1 (BCC) or Cyclin A2 (AC) (Figure 1A). CDK1 was shown to be sufficient to drive cell cycle progression in the absence of the interphase CDKs *in vivo* (Santamaría et al., 2007). Therefore, we used centrifugal elutriation to enrich for cells in the G2&M phases of the cell cycle (fractions 11 and 12 in Supplementary Figure 1A). In these cells, targets of interphase CDKs (CDK2/4/6) will be phosphorylated and protein substrates critical to the essential role of CDK1 in mitotic entry are expressed. Increased phosphorylation on SPXK motifs was observed after reaction with complexes containing active CDK1, but not a kinase dead CDK1 mutant (CDK1^D146N^, KD) (Figure 1B). Phosphorylation is a readout for two molecular events: an interaction between the recombinant CDK1 added and the substrate, and catalytic phosphotransfer. Because cells are fixed, this phosphorylation is direct and not subject to feedback (e.g., dephosphorylation by phosphatases). We then prepared peptide digests from the phosphorylated cells using an in-cell digest (Kelly et al., 2022). Peptide digests were labelled with 16-plex tandem mass tags (TMTs), enriched for phosphorylated peptides, fractionated offline by HPLC prior to LC-MS/MS analysis (Figure 1A). 3,377 phosphorylation sites, representing 2,493 proteins, were increased by 2-fold or more compared to KD. Motif enrichment analysis of significantly changing phosphorylation sites revealed a known consensus sequence for CDK1 (TPXK) (Figure 1C). In contrast, sites significantly increased after treatment with Aurora B are enriched in an RRX[ST] motif (Figure 1D). Our results recapitulate consensus motifs obtained using cell-based assays, indicating that sites identified in fixed cell assays are specific to the kinase added (Holt et al., 2009; Kettenbach et al., 2011).

**Figure 1.**
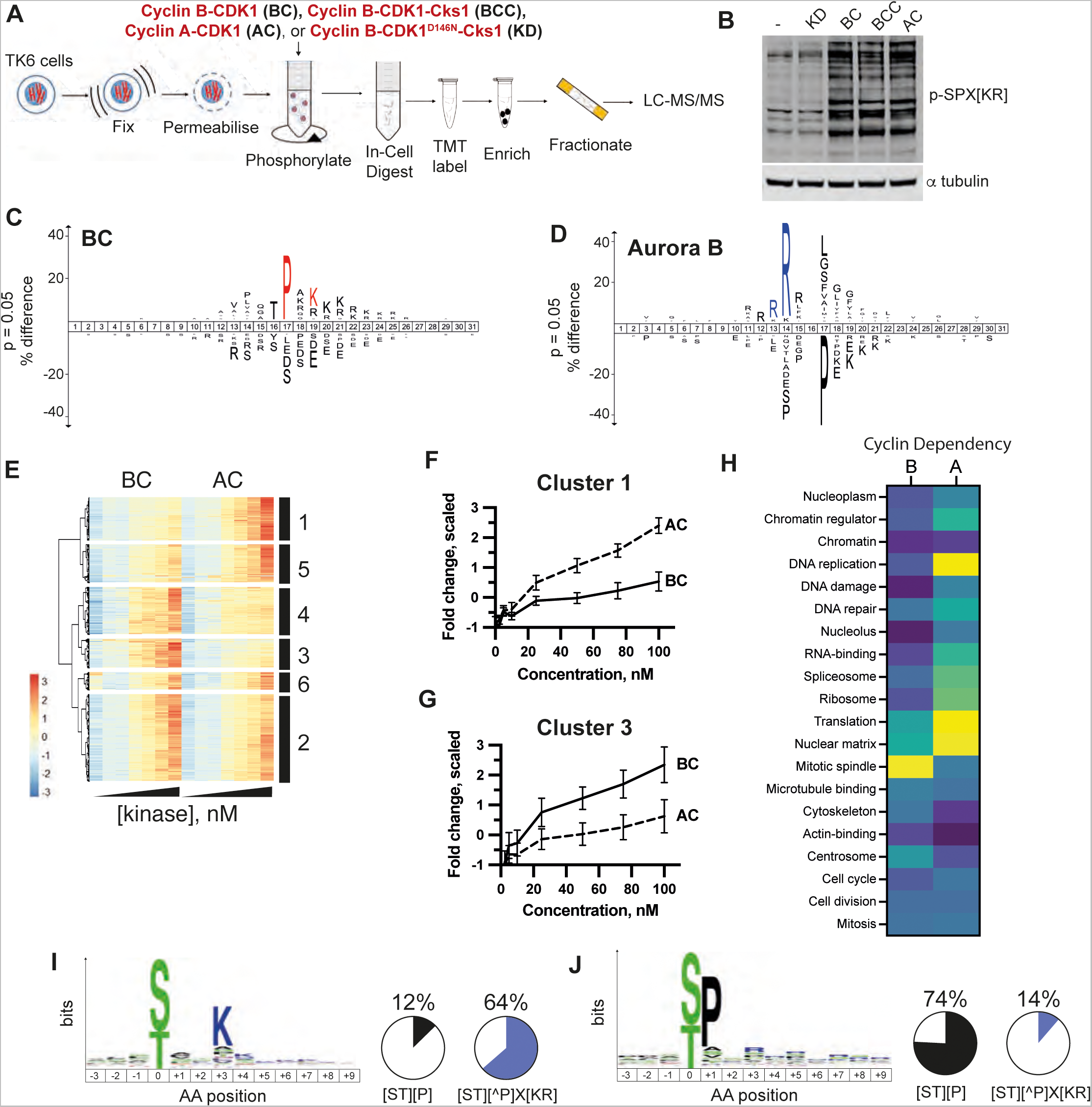
Cyclin A2 promotes non-Proline directed CDK1 phosphorylation and modulates substrate specificity *in vitro*. (A) Scheme of the *in vitro* kinase assay on fixed cells, followed by MS-based phosphoproteomics. Cyclin B-CDK1 (BC), Cyclin B-CDK1-Cks1 (BCC), Cyclin A-CDK1 (AC), and a kinase dead mutant of CDK1, Cyclin B-CDK1^D146N^-Cks1 (KD) were compared. (B) Western blot of lysates from fixed cells phosphorylated using the assay described in (A) with anti-Phospho-SPX[KR] motif antibody, which also cross-reacts with phosphorylated MAPK motifs, i.e., PXpSP. (C) Motif enrichment analysis of fixed G2&M cells phosphorylated by either BC or BCC CDK1 complexes. Amino acids shown on top and bottom are enriched and under-represented, respectively, in kinase phosphorylated sites, compared to background. The amino acid in position 0 represents the phospho-acceptor residue. (D) Motif enrichment analysis of fixed G2&M cells phosphorylated by Aurora B. (E) Fixed G2&M cells were titrated with increasing concentration of either AC or BC, and subjected to phosphoproteomic analysis. Colour indicates the scaled fold change relative to KD. (F, G) The mean fold change plotted against the concentration of AC and BC for cluster 1 (F) or cluster 3 (G). Error bars represent the standard deviation. (H) Heatmap showing fold enrichment for selected gene ontology (GO) terms and UniProt keywords comparing Cyclin A- vs Cyclin B-dependent phosphorylation sites. (I, J) Motif enrichment analysis of sites from cluster 1 (I) or cluster 3 (J). Proportion of sites matching +1P (proline-directed, black) or S[^P]X[KR] (blue) motifs, where ^P denotes any amino acid besides P and X is any amino acid.

### Cyclin A2 shifts the substrate specificity of CDK1 and promotes the phosphorylation of non-Proline sites in vitro

To investigate how the quality and quantity of CDK1 affects substrate choice, we used the fixed cell assays described above to measure CDK1 substrate phosphorylation proteome-wide comparing AC and BC titrated from 2.5 nM to 100 nM recombinant kinase. An upper limit of 100 nM was chosen based on estimated copies of Cyclin B1 and CDK1 detected in G2&M-phase leukemic cells (Ly et al., 2014; Wiśniewski et al., 2014). 27,084 phosphorylation sites were detected, of which 5,113 sites changed by 2-fold or more compared to KD-treated cells (Supplementary Table 1). This dataset allowed us to investigate how increasing CDK1 activity quantity affects substrate choice by measuring concentration dependent changes in substrate phosphorylation. On the other hand, with the same dataset, we can investigate how the quality of CDK1 activity (i.e., the different Cyclin subunit) impacts substrate choice.

To assess the relative differences between Cyclin A and Cyclin B phosphorylation for the same site, fold changes were scaled (see Methods) so that the means and standard deviations for all sites were identical. Scaled fold changes were then normalized for differences in kinase activity between recombinant kinase preparations (Supplementary Figure 1B). Histone H1 protein has been used extensively to assess specific activity of CDK (e.g., Loog and Morgan, 2005). Normalized, concentration-dependent phosphorylation of histone H1 is indistinguishable between AC and BC (Supplementary Figure 1C). Thus, any differences between AC and BC observed in the subsequent analysis are not due to variation in kinase activity, including specific activity for histone H1, and instead, are due to qualitative differences conferred by the Cyclin subunit.

Hierarchal clustering of these data to identify groups of phosphorylation sites that exhibited similar patterns of phosphorylation. This clustering identified six major clusters of phosphorylation sites (Figure 1E). Sites in cluster 1 were phosphorylated to a higher extent with AC compared to BC and therefore Cyclin A-dependent (Figure 1F). Vice versa, sites in cluster 3 were Cyclin B-dependent (Figure 1G). The observation of equivalently sized clusters showing either AC, or BC dependence, supports the idea that the phosphorylation patterns observed are unlikely due to differences in the specific activity of the purified kinases.

Are there characteristics that distinguish Cyclin A- versus Cyclin B- dependent phosphorylation sites and substrate proteins? To address this question, we compared clusters 1 and 3 because they show the most extreme differences in cyclin subunit dependence. First, we assessed the frequency of annotated biological function, subcellular localization gene ontology terms and UniProt keywords to see if any annotations were differentially enriched (Figure 1H). Cyclin A-dependent protein substrates were differentially enriched in proteins with functions in DNA replication, protein translation and proteins localized to the nuclear matrix. In contrast, Cyclin B-dependent protein substrates were enriched in proteins localized to the mitotic spindle and centrosomes. Cyclin A- and Cyclin B-dependent substrates were equally enriched in proteins with functions in mitosis and cell division (Figure 1H, bottom). No significant difference was observed between Cyclin A- versus Cyclin B-dependent substrates in whether they contain 1 or more RXL motifs in the same disordered region as the phosphorylation site (Fisher’s exact test, p > 0.05).

Cyclin A-dependent sites show a remarkable depletion of Proline in the +1 position, with only 12% sites being proline-directed (Figure 1I). In contrast, Cyclin B-dependent sites show a strong +1 Proline (+1P) preference, with 74% sites being proline-directed. (Figures 1J). Cyclin A-dependent sites contain a prominent enrichment of Lysine at the +3 position (Figure 1I). 64% of Cyclin A-dependent sites meet the S[^P]XK motif (Figure 1I). In contrast, 14% of Cyclin B-dependent sites meet this S[^P]XK motif (Figure 1J).

We conclude that the Cyclin subunit has a major role in subcellular and substrate targeting. Cyclin A increases the frequency of non-proline directed CDK1 phosphorylation compared to Cyclin B, and these sites are enriched in S[^P]XK sites.

### Quantitative increases in CDK1 activity alters substrate specificity in vitro

Because phosphorylation does not saturate in the range of kinase concentrations tested (Figure 1E), using scaled (relative) fold changes, we cannot assess whether sites differ in concentration dependence. For many sites, phosphorylation is not detected in KD-treated cells, therefore making exact fold changes challenging to accurately estimate. Therefore, to facilitate comparison of concentration dependence between sites, an arbitrary fold-change cap was used to enforce phosphorylation saturation *in silico*. A cap of 6.5 was chosen because this was the mean fold change for phosphorylation sites increased in mitotic versus G2 cells. This approach enables us to determine if, and at what concentration, the phosphorylation effectively reaches the mean fold-change observed in the G2 to M transition in cells.

Using this strategy, we identified seven clusters of phosphorylation sites that differed in BC concentration-dependence (Figures 2A and B, Supplementary Table 2). Cluster 1 shows fold changes of 6.5 or more at 2.5 nM, the lowest concentration tested. At the other extreme, sites in cluster 7 do not reach fold changes of 6.5 or more at 100 nM and follow a linear response to kinase concentration. Proteins in these clusters were then subjected to functional gene annotation enrichment analysis. Figure 2C shows selected functional gene ontology (GO) annotations that show a significant (FDR < 0.01) in one or more clusters. Cluster 1 shows no major differential enrichment. Interestingly, Cluster 2 shows an enrichment in proteins associated with protein translation, including ribosomal proteins. Sites with moderate sensitivity to CDK1 activity (cluster 4), are significantly enriched in proteins involved in DNA repair and DNA replication. Sites with lower CDK1 sensitivity (cluster 6) are enriched in proteins involved in the organization of the mitotic spindle, chromosome segregation and nuclear envelope disassembly. For example, proteins in cluster 6 show a 7.6-fold enrichment in mitotic spindle assembly proteins, which is three times higher than cluster 1. Similar trends in clustering and functional enrichment are observed with AC (Supplementary Figures 1D, E). Individual CDK1 substrate proteins may have sites belonging to multiple clusters. Indeed, 943 out of 2,550 CDK1 substrate proteins have phosphorylation sites with differing sensitivities to CDK1 phosphorylation. CDK1 phosphorylation sites on these substrates represent the majority of the phosphorylation sites (64%, 3,288 / 5,113). We conclude that quantitative changes in the activities of Cyclin A-CDK1 and Cyclin B-CDK1 alter substrate specificity in a similar concentration-dependent manner, with many substrates having multiple CDK1 sites with differing sensitivity to kinase activity.

**Figure 2.**
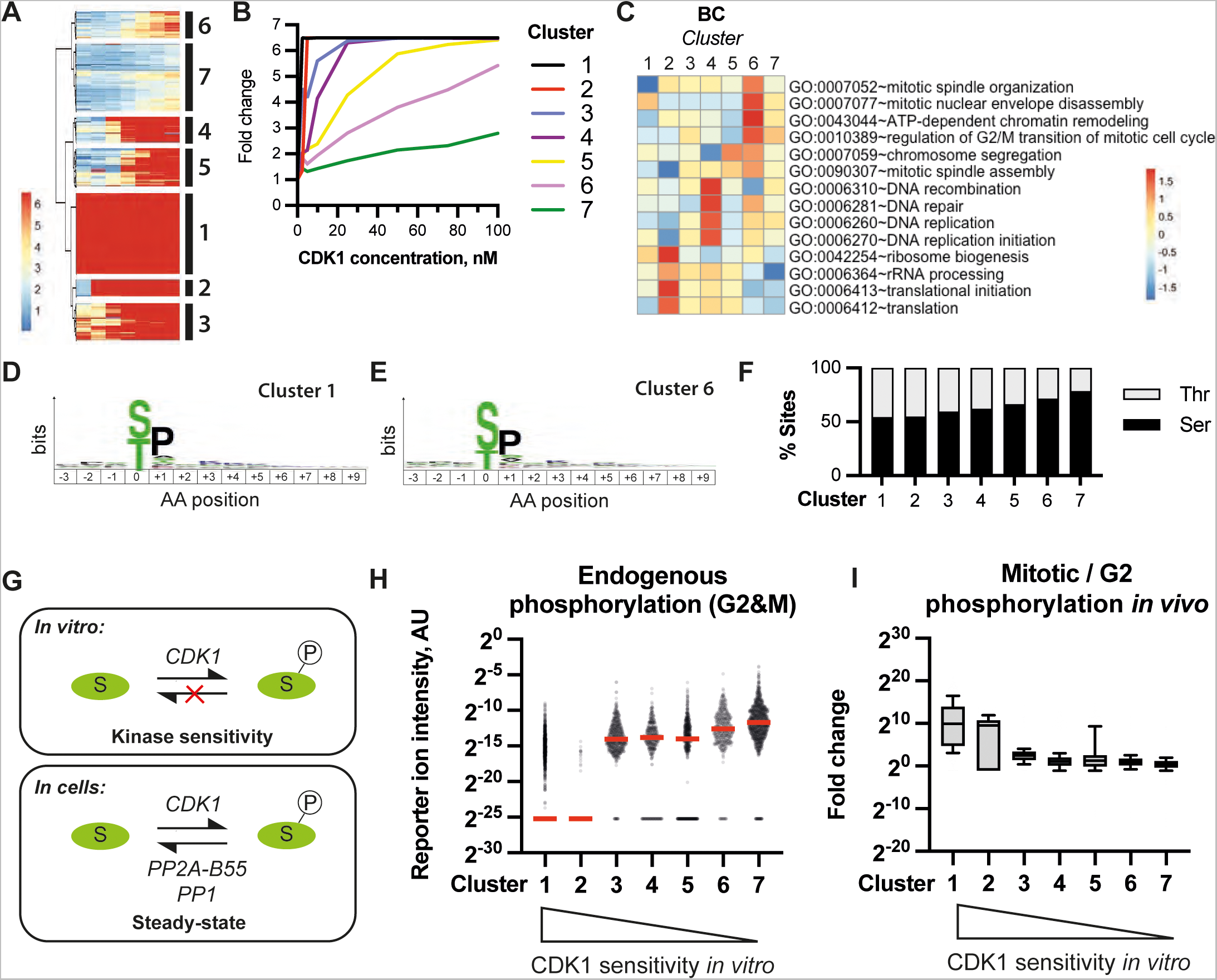
Quantitative increases in CDK1 activity alters substrate specificity *in vitro*. (A) G2&M cells were titrated with increasing concentrations of KD or BC and subjected to phosphoproteomic analysis. Heat map showing the capped fold changes against KD. Hierarchical clustering identified seven clusters. (B) Mean capped fold-change versus CDK1 concentration for each cluster. (C) Scaled fold enrichment for selected enriched gene ontology terms. (D, E) Motif analysis of sites in Cluster 1 (D) and Cluster 6 (E). (F) Proportion of serine and threonine phosphoacceptor residues per cluster. (G) Model for the readout of phosphorylation in fixed cells (kinase sensitivity) versus in viable cells (steady- state levels, which is subject to kinase-phosphatase antagonism). (H) Normalized phosphopeptide intensities detected in KD-treated G2&M cells (i.e., the endogenous phosphorylation in cells) per cluster. Bars indicate medians. (I) Fold-change comparing mitotic (SLTC-arrested) versus G2&M phase cells per cluster. Bars indicate medians, and box and whiskers show 75^th^/25^th^ and 90^th^/10^th^ percentiles, respectively.

We next assessed if clusters differed in amino acid sequence proximal to the phosphoacceptor residue. Low and high CDK1 sensitivity clusters showed a similar preference for +1 Proline (Figures 2D, 2E). Similar results were obtained using Cyclin A-CDK1 (Supplementary Figure 1F). Interestingly, high CDK1 sensitivity sites in cluster 1 have a high proportion of threonine phosphoacceptor sites (∼50%, Figure 2F). In contrast, the proportion of threonine sites (over serine) is 36% and 23% for CDK1 sites and all detected sites, respectively. Indeed, the proportion of threonine phosphoacceptor residues is negatively correlated with CDK1 concentration dependence (Figure 2F). These results suggest that threonine residues are a major target of CDK1 activity in the G2 to M transition.

In the fixed cell assay, substrate phosphorylation will be determined solely by the forward kinase reaction. However, in cells, substrate phosphorylation will be subject to antagonism by cellular protein phosphatases, including PP2A-B55 and PP1, as illustrated in Figure 2G. PP2A:B55 phosphatase has been shown to prefer dephosphorylating phosphothreonine residues with a +1 Proline and is active in interphase (Cundell et al., 2016). Could targeted dephosphorylation by PP2A:B55 *in vivo* lead to lower occupancy of TP sites in fixed G2 cells and therefore higher CDK1 sensitivity *in vitro*? To address this question, we examined the endogenous abundance of phosphorylation in G2&M cells, grouped into phosphorylation sites determined by CDK1 sensitivity by *in vitro* (Figure 2H). Cluster 1 has the highest proportion of threonine sites and the lowest level of endogenous phosphorylation. Indeed, there is a negative correlation between CDK1 sensitivity *in vitro* and the median level of endogenous phosphorylation (Figure 2H). As will be elaborated in the discussion, kinase reactions with G2 cells pre-treated with lambda phosphatase would ensure that all sites have identical phosphorylation stoichiometry at the start of the kinase reaction. Within a cluster, however, individual sites range widely in levels of endogenous phosphorylation (Figure 2H), suggesting that the sensitivity towards CDK1 *in vitro* is unlikely to be driven exclusively by endogenous occupancy. Does CDK1 sensitivity *in vitro* reflect *in vivo*? To test this, we plotted endogenous phosphorylation fold changes comparing mitotic and G2 cells because in mitosis, PP2A-B55 is inactivated and CDK1 activity is high. Indeed, sites showing the highest sensitivity to CDK1 *in vitro* show the highest fold changes in mitosis (Figure 2I). These data support a model whereby phosphatase inactivation is the crucial determinant of CDK1 substrate phosphorylation (Castilho et al., 2009; Krasinska et al., 2011; Mochida et al., 2009; Vigneron et al., 2009).

### Cks1 promotes the phosphorylation of non-Proline sites by CDK1 in vitro

To assess the role of Cks1 in Cyclin B1-CDK1 interaction with cellular substrates, we titrated fixed G2&M-enriched TK6 cells with increasing concentrations of either BC or BCC, performed in biological duplicate. Out of 24,139 sites identified (Supplementary Table 3), 3,377 sites showed a 2-fold or more increase in phosphorylation upon addition of active CDK1. Note that the data for BC is identical to that shown in Figure 1. Fold changes were normalized for differences in CDK1 activity, as above (Supplementary Figure 2A). Individual sites differed in CDK1 concentration dependence (Figure 3A), and some sites were phosphorylated to higher extent with BCC compared with BC, and vice versa. Phosphorylation sites were separated into four clusters by hierarchal clustering. Phosphorylation sites in cluster 2 show strong enhancement by the Cks1 subunit (Figure 3B). The remaining sites were either phosphorylated to a greater extent by BC (cluster 4, Figure 3C, ‘Cks1-inhibited cluster’), or equally phosphorylated by BC and BCC (cluster 3). Interestingly, there is a small set of phosphorylation sites that are highly phosphorylated by BCC at 100 nM kinase (cluster 1).

**Figure 3.**
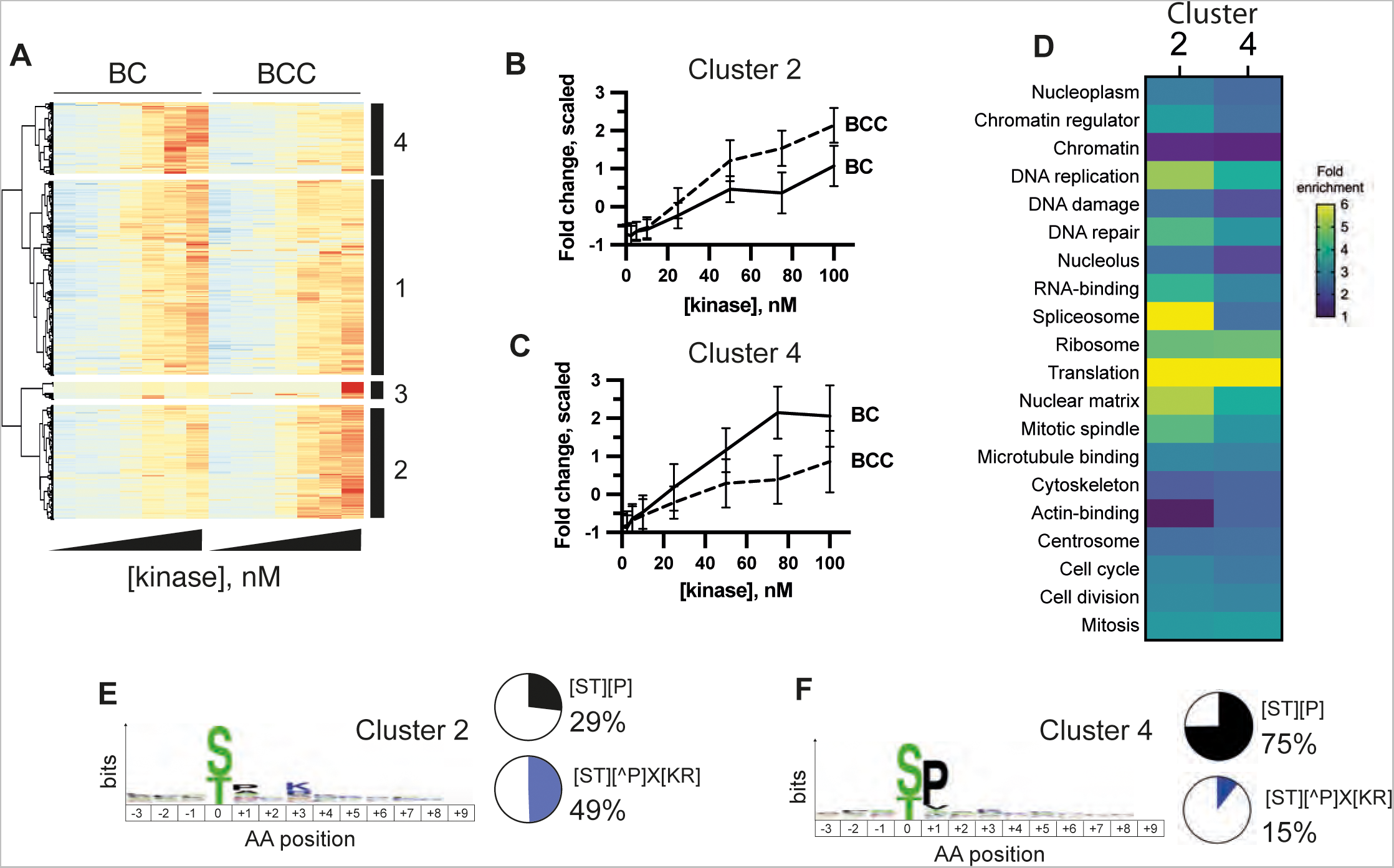
Cks1 promotes widespread phosphorylation of non-Proline directed sites by human Cyclin B-CDK1 *in vitro*. (A) G2&M cells were titrated with increasing concentrations of KD, BC or BCC and subjected to phosphoproteomic analysis. Heat map showing the scaled fold changes against KD. Hierarchical clustering was used to identify Cks1- dependent (Cluster 2) and Cks1-independent (Cluster 4) phosphorylation sites. (B, C) Line graphs showing mean fold change for Clusters 2 (B) and 4 (C). Error bars represent the standard deviation. (D) Heatmap showing fold enrichment for selected gene ontology (GO) terms and UniProt keywords comparing Cks1-dependent vs Cks1-independent phosphorylation sites. (E, F) Motif analysis of the sites in Clusters 2 (E) and 4 (F) using WebLogo. Pie charts show the proportions of sites in the cluster with the indicated motif, e.g., +1P (black), or S[^P]X[KR] (blue), where ^P denotes any amino acid besides P and X is any amino acid.

How do Cks1-enhanced sites differ from Cks1-inhibited ones? A functional enrichment analysis was performed comparing clusters 2 and 4. In general, the differences seen between Cks1-enhanced and Cks1-inhibited substrates (Figure 3D) are less than those observed between Cyclin A-CDK1 and Cyclin B-CDK1. Splicing factors, however, are a striking exception, being highly enriched in Cks1-enhanced substrates. Enrichment is also slightly higher in Cks1-enhanced substrates for proteins localized to the nuclear matrix and with functions in DNA replication. These results support a model whereby BCC increases phosphorylation of a subset of BC substrates.

We next tested if the local amino acid sequence around the phospho-acceptor site in the Cks1-enhanced cluster (cluster 2) is significantly different from a Cks1-inhibited cluster (cluster 4). As shown in Figure 3E, Cks1-enhanced sites are depleted of Proline in the +1 position (+1P). Strikingly, only 27% of Cks1-enhanced sites had a +1P, in contrast to Cks1-inhibited sites, in which 91% had a +1P (Figure 3F). There is a slight preference for a lysine in the +3 position (+3K), but less than Cyclin A-dependent sites (Figure 1I). 49% of Cks1-enhanced sites meet a S[^P]XK motif. In contrast, only 3% of Cks1-enhanced sites meet this S[^P]XK consensus (Figure 3F). We conclude that Cks1 has a proteome-wide role in promoting CDK1 phosphorylation of non-proline directed sites *in vitro*.

### Cks1 enhances CDK1 primed-multisite phosphorylation of substrate proteins in vitro

Cks1 has been shown to enhance substrate multisite phosphorylation of the protein substrate, *S. cerevisiae* Sic1, by phospho-dependent docking to a priming site (Kõivomägi et al., 2013; Köivomägi et al., 2011). Cks1 was proposed to act as a molecular ruler, promoting the phosphorylation of a second, lower affinity site 12-15 amino acids C-terminal to the priming site (Figure 4A). To what extent does phosphate docking play a role in CDK1 phosphorylation in human cells? And is this role of Cks1 proteome-wide?

**Figure 4.**
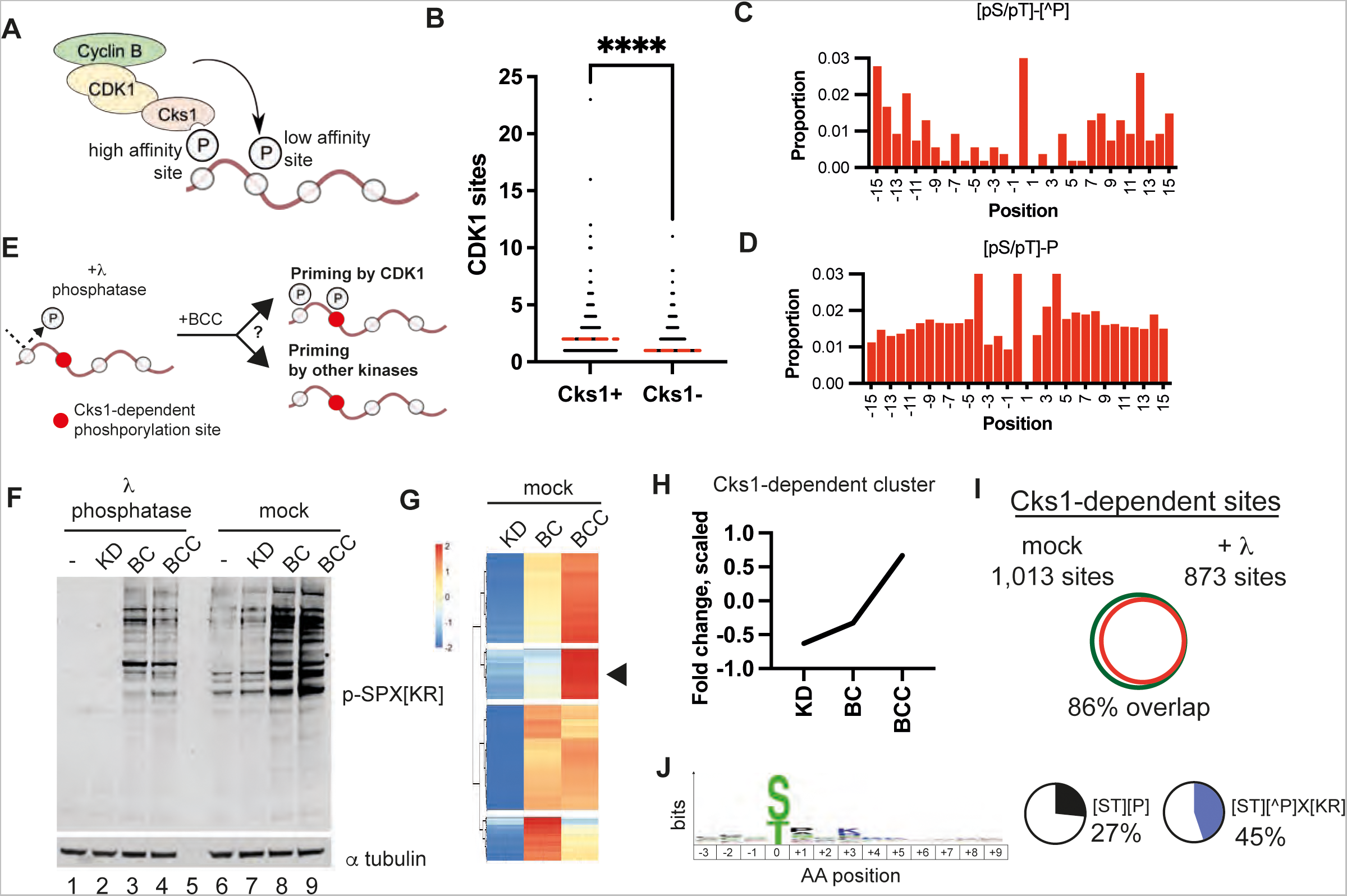
Cks1 promotes multisite phosphorylation of CDK1 protein substrates. (A) Model for Cks1 function in CDK1 substrate phosphorylation. (B) Dot plot representing the number of sites (*y-axis*) phosphorylated on each protein substrate (dot). Only proteins unique to each cluster are shown. \*\*\*\**p*<0.0001 from a student’s *t* test; dotted lines represent the median. (C, D) The frequency of additional phosphorylation sites detected by phosphoproteomics within 15 residues of the phosphoacceptor for Cks1-dependent sites lacking the +1P (C), and CDK1 sites with a +1P (D). (E) Experimental design to investigate priming kinases for Cks1. (F) Western blot with anti-phospho-SPX[KR] motif antibody of lysates from fixed cells phosphorylated *in vitro* pre-treated either with λ Phosphatase or mock. This antibody cross-reacts with phosphorylated MAPK motifs, i.e. PXpSP. (G, H) Heatmap and line graph showing the identification of Cks1-dependent phosphorylation sites in mock-treated cells. (I) Overlap between sites that show Cks1-dependence in mock-versus λ Phosphatase-treated cells. (J) Motif enrichment analysis of overlapping sites. Pie charts show the proportions of sites in the cluster with the indicated motif, e.g., +1P (black), or S[^P]X[KR] (blue), where ^P denotes any amino acid besides P and X is any amino acid. Reproducible sites from two biological repeats are shown.

We reasoned that if Cks1 promoted multisite phosphorylation, then Cks1-enhanced protein substrates should show on average, a higher number of sites phosphorylated by CDK *in vitro* than Cks1-inhibited substrates. Indeed, protein substrates were exclusively Cks1-enhanced had on average two CDK1 phosphorylation sites per protein (median), compared with one site per protein for Cks1-inhibited substrates (Figure 4B). Many proteins had multiple sites, some that are Cks1-enhanced and others that are Cks1-inhibited. For example, on the protein Ki- 67, 49 sites are phosphorylated by CDK1 in total, of which 24 are Cks1-enhanced. Unlike Cks1, the Cyclin subunit does not alter multisite substrate phosphorylation (Supplementary Figure 2B).

Next, we examined the distribution of secondary phosphorylation sites surrounding a Cks1- dependent phosphoacceptor site. We reasoned if Cks1 acts as a molecular ruler, we should observe ‘hot spots’ of secondary phosphorylation where priming phosphorylation is preferred. In support of this model, secondary phosphorylation sites are enhanced at positions -15 and +12 for BCC-dependent non-proline sites (Figure 4C). Interestingly, proline-directed CDK1 sites overall do not show this behavior (Figure 4D). The enhancement is seen for both serine and threonine phosphoacceptor residues, and there is no difference in pattern if the secondary (putative priming phosphorylation) is restricted to either serine, or threonine. Interestingly, there is a depletion of secondary phosphorylation sites from positions -8 to +6 for BCC- dependent non-proline sites compared with proline-directed sites. These results support a model whereby Cks1 docks onto a proximal priming site to facilitate multisite phosphorylation, frequently at non-proline directed sites.

The identity of the major priming kinase for Cks1 is unknown. In *S. cerevisiae*, CDK-Cks complexes can self-prime to phosphorylate a substrate in a processive manner (Kõivomägi et al., 2013). Our data suggest that docking can occur either N-terminal or C-terminal to the phosphoacceptor residue, and that the priming phosphorylation can be either serine or threonine. However, phosphoproteomic analysis is not at saturation, and the bioinformatic analysis above cannot distinguish multiple proteoforms of the same protein that differ in phosphorylation (e.g., two proteoforms each phosphorylated at different sites, or a single proteoform phosphorylated at both sites).

Therefore, to directly investigate priming, we designed an experiment applying sequential phosphatase and kinase reactions on fixed cells. We reasoned that removal of all endogenous phosphorylation in G2 cells would eliminate priming by all kinases except for the one added (CDK1). By using a proteome-wide approach, we can identify which Cks1-dependent sites are dependent on priming by CDK1, or by other kinases (Figure 4E).

TK6 cells were treated with λ Phosphatase, followed by BCC. Phosphatase pre-treatment eliminated endogenous SPX[KR] phosphorylation (Figure 4F, lanes 1 and 6), and SPX[KR] phosphorylation is increased after treatment with BC or BCC (Figure 4F, e.g., compare lanes 2 and 3). Reactions were then analyzed by LC-MS/MS as shown in Figure 1A. 23,911 sites were identified, of which 5,819 sites were phosphorylated by CDK1 (G2&M cells, Supplementary Table 4). Sites phosphorylated by CDK1 in mock treated cells were clustered (Figure 4G) to identify Cks1-enhanced sites (Figure 4G, arrow, and Figure 4H). We then asked if these sites were phosphorylated by CDK1 in λ phosphatase-treated cells. Of the 1,013 Cks1- enhanced sites phosphorylated in mock-treated cells, 873 sites showed a 2-fold or higher change after addition of BCC to λ phosphatase-treated cells compared to KD (Figure 4I). This result demonstrates that CDK1 has priming activity for most Cks1-dependent sites (86%). The remaining 140 Cks1-dependent sites cannot be primed by the CDK1 complexes tested, and are likely to be primed by other kinases, or else mediated by phosphorylation-independent docking interactions. Interestingly, these experiments, which were carried in biological duplicate separately from those shown in Fig. 3E, also show that Cks1-dependent sites are depleted of the +1P and are instead enriched for a +3K (Figure 4J).

Taken together, our results here have shown that a subset of CDK1 phosphorylation sites lack the +1 Proline consensus and that phosphorylation of these non-proline directed sites by BCC is enhanced by the phospho-adaptor protein, Cks1. Cks1-dependent phosphorylation, which is primed by CDK1 itself *in vitro* promotes the multisite phosphorylation of protein substrates.

### Cks1- and Cyclin A- dependent phosphorylation sites are cell cycle regulated

A crucial question is whether CDK1 phosphorylation observed *in vitro* is also observed in living cells. The primary function of CDK activity is to drive cell cycle progression via ordered phosphorylation of substrates, leading to increasing phosphorylation occupancy across the cell cycle with maximal occupancies observed in mitosis (Swaffer et al., 2016). Therefore, we hypothesized that if *in vitro* sites are physiologically relevant, they should also be cell cycle regulated (CCR) *in vivo*. We designed an experiment where cells enriched in different cell cycle stages were directly compared against fixed cells phosphorylated *in vitro* by CDK1 (Figure 5A). These samples were analyzed together in a single quantitative TMT phosphoproteomics batch, which minimizes missing values. This internally controlled, relative quantitation approach enables straightforward and comprehensive comparison between *in vitro* and *in vivo*.

**Figure 5.**
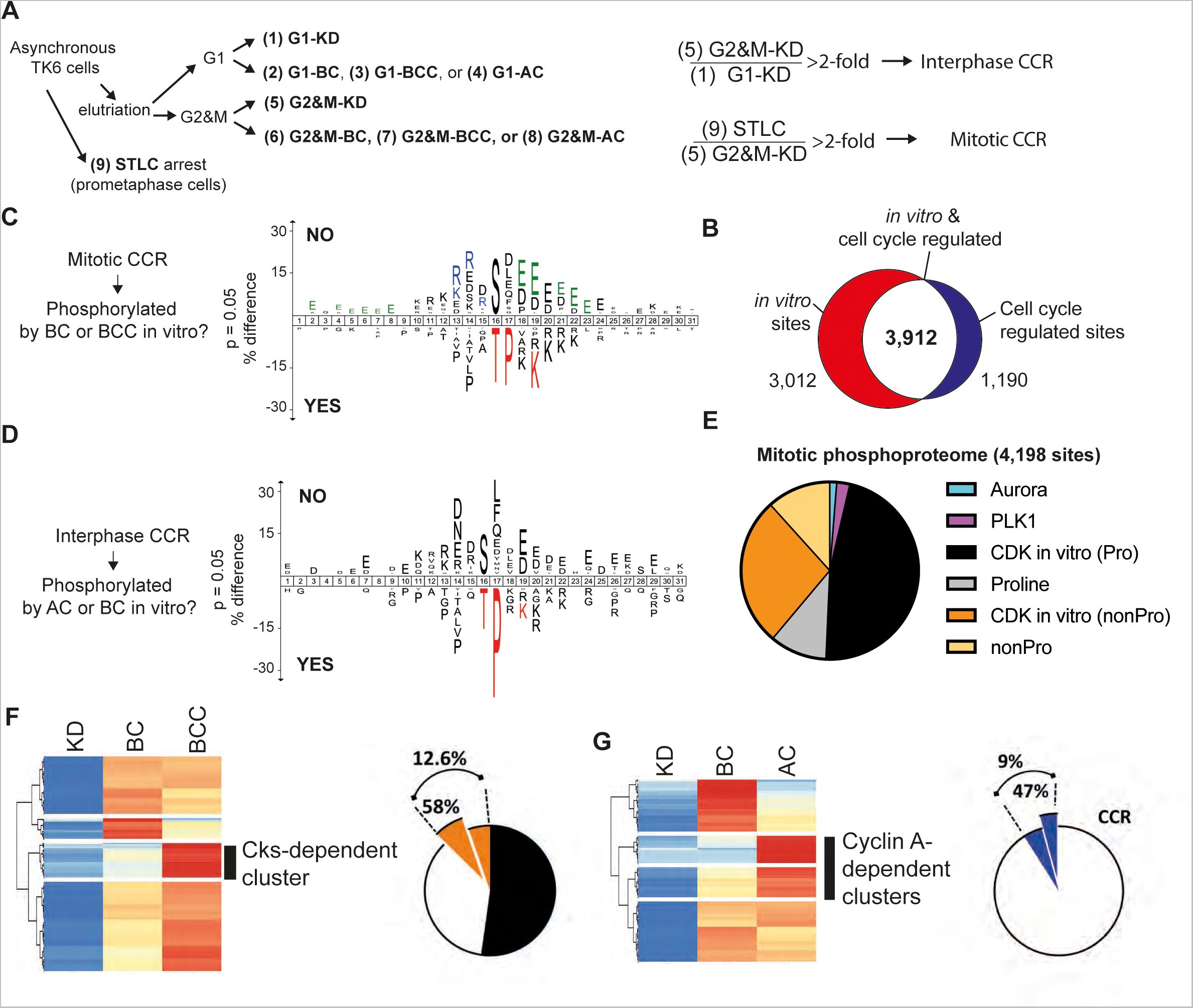
Non-proline directed CDK phosphorylation sites *in vitro* are cell cycle regulated in viable cells. (A) Design of experiment to identify both cell cycle regulated phosphorylation sites and *in vitro* CDK phosphorylated sites. All samples were combined into a single TMT analysis to minimize missing values. (B) Overlap between sites phosphorylated *in vitro* by AC, BC, or BCC, and the cell cycle regulated phosphoproteome. (C) Enriched motifs for mitotic CCR sites, that were either not phosphorylated by CDK1 *in vitro* (top) phosphorylated by CDK1 *in vitro* (bottom). (D) Motif enrichment analysis of interphase CCR sites that did not overlap with CDK1 *in vitro* targets (top) in comparison to that of those that overlapped (bottom). (E) The proportion of phosphorylation sites in the mitotic regulated phosphoproteome that can be explained by consensus and direct CDK1 phosphorylation *in vitro*. (F) Heatmap showing mitotic CCR phosphorylation sites that are Cks1-dependent *in vitro* (orange), including non-proline directed sites (exploded pie slice). (G) Heatmap showing interphase CCR phosphorylation sites that are Cyclin A-dependent *in vitro* (blue), including sites that meet the S[^P]XK consensus (exploded pie slice).

G1- and G2&M populations were collected using centrifugal elutriation (Supplementary Figure 3A). 9% of cells in the G2&M-enriched fraction were mitotic, judging by Histone H3 S10 phosphorylation (H3pS10). In parallel, cells were arrested in prometaphase using STLC. Fixed G1 and G2&M-enriched cells were subjected to kinase assays using AC, BC, BCC, and KD, as above. 23,911 sites were detected in total (Supplementary Tables 5 and 6), representing 5242 protein substrates. 1,514 sites showed 2-fold or more phosphorylation in the G2&M-enriched sample compared to G1 cells. These sites were deemed interphase CCR (Figure 5A). Similarly, 4,198 sites showed 2-fold or more phosphorylation in STLC-arrested cells compared to the G2&M population and deemed to be mitotic CCR (Figure 5A). In total, there were 5,102 sites either mitotic or interphase CCR. 6,924 sites showed 2-fold or more phosphorylation *in vitro* after incubation with one of the three active CDK1 complexes compared to samples treated with KD. Of these, 3,912 sites overlapped with either interphase or mitotic CCR sites (Figure 5B). SPXK phosphorylation in STLC-arrested mitotic cells is more intense than G2&M-enriched cells subjected to phosphorylation by CDK1 *in vitro*, demonstrating that the phosphorylation occupancy achieved *in vitro* is at levels below the maximum achieved physiologically (Supplementary Figure 3B). These data demonstrate the fixed cell kinase assays phosphorylate sites to physiologically relevant levels, and that a majority of *in vitro* CDK1 sites (56%) are phosphorylated in a cell cycle regulated manner.

Sequence analysis of mitotic CCR sites not phosphorylated by CDK1 *in vitro* show a significant enrichment in motifs consistent with Aurora A/B and acidophilic kinases, including Plk1, Polo-like Kinases 2/3 and Casein Kinase 2 (CK2) (Figure 5C, top). Interphase CCR sites not phosphorylated by CDK1 *in vitro* show enrichment in acidophilic kinases, but no enrichment of Aurora A/B motifs. Non-CDK1 interphase CCR sites have a prominent enrichment of an acidic residue in the -2 position, consistent with a relaxed Plk1 consensus motif (Figure 5D, top). Both interphase and mitotic sites phosphorylated by CDK1 *in vitro* show an enrichment of the classic CDK consensus sequence (Figures 4C and 4D, bottom), which is virtually identical to a similar analysis of all *in vitro* CDK1 sites, including non-CCR sites (Figure 5C).

Next, we asked whether non-proline directed sites phosphorylated by CDK1 *in vitro* are CCR in living cells (Figure 5E). Of the 4,198 mitotic CCR sites, 2,413 sites (57%) met a +1P minimum CDK consensus. Of these +1P CCR sites, 1,975 (82%) were phosphorylated by CDK1 *in vitro*. Only 100 and 53 of the mitotic CCR sites meet the strict consensus motifs for Polo-like Kinase 1 (Plk1) and Aurora Kinases ([KR][KR]X[ST]), respectively. The remaining 1,632 non-proline directed mitotic CCR sites, which do not meet any of the motifs described above, constituted 39% of the total mitotic CCR phosphoproteome. Strikingly, 1,142 of these non-proline directed mitotic CCR sites, representing 70% of phosphorylation sites with no predicted upstream cell cycle kinase, are phosphorylated by CDK1 *in vitro*. We therefore conclude that the majority of non-proline directed mitotic CCR phosphorylation sites can be directly phosphorylated by CDK1. Taken together, our data show that 3,117 out of 4,198, or 74%, of the mitotic CCR phosphoproteome can be phosphorylated directly by CDK1 (proline and non-proline directed) (Figure 5E).

We then examined if there is evidence that Cks1-enhanced and Cyclin A-dependent phosphorylation sites are CCR. Indeed, Cks1-enhanced sites *in vitro* (Figure 5F and Supplementary Table 5) constitute 12.6% of the mitotic CCR phosphoproteome, of which 58% are non-proline directed. These Cks1-dependent, mitotic CCR sites show depletion of a +1P (Supplementary Figure 3C). We reasoned that because Cyclin A2 is degraded during an extended prometaphase arrest (Supplementary Figure 3B), Cyclin A2-dependent sites will be subject to attrition in the STLC-arrested sample. CDK1 can phosphorylate substrates of interphase CDKs (CDK2/4/6). Therefore, to test if Cyclin A-dependent phosphorylation sites are also CCR, we performed the fixed cell kinase assay on interphase cells. G1 cells were phosphorylated with either BC or AC *in vitro*. Hierarchical clustering identified a group of sites that were phosphorylated *in vitro* in a Cyclin A-dependent manner (Figure 5G and Supplementary Table 6). These sites represented 9% of the interphase CCR sites and 42% of these sites matched the [ST][^P]XK motif (Figure 5G). Sequence motif analysis of Cyclin A-dependent, interphase CCR sites show a depletion of a +1P and an enrichment of a +3K (Supplementary Figure 3D).

### A [ST][^P]XK sequence motif is enriched among non-proline directed CDK1 sites

Our data show that both Cks1 and Cyclin A increase the frequency of non-proline directed sites. Unlike Cyclin A, Cks1 is not targeted for degradation by APC/C-Cdc20 in early prometaphase, and is found in cells in complex with Cyclin B-CDK1. Interestingly, a pool of Cyclin B-CDK1 is localized to the kinetochore corona and plays a role in spindle assembly checkpoint signaling (Allan et al., 2020). We wondered if there was overlap between Cks1- and Cyclin A-dependent sites because phosphorylation of these sites could be ‘handed over’ from Cyclin A-CDK1 (with, or without Cks1) to Cyclin B-CDK1-Cks1 during prometaphase after Cyclin A is degraded. The overlap between sites reproducibly dependent on Cyclin A (red, N = 2) and Cks1 (blue, N = 2) is shown in Figure 6A (Supplementary Table 7). 44 sites on 41 proteins were in common. These included sites on Ki-67, MCAK (KIF2C) and Hec1 (NDC80), which all have functions in regulating chromosome segregation. The [ST][^P]X[K] motif that was observed individually for Cks1- and Cyclin A-dependent sites is strongly enriched in these overlapping sites (Figure 6B) and are found in 39 out of the 44 sites.

**Figure 6.**
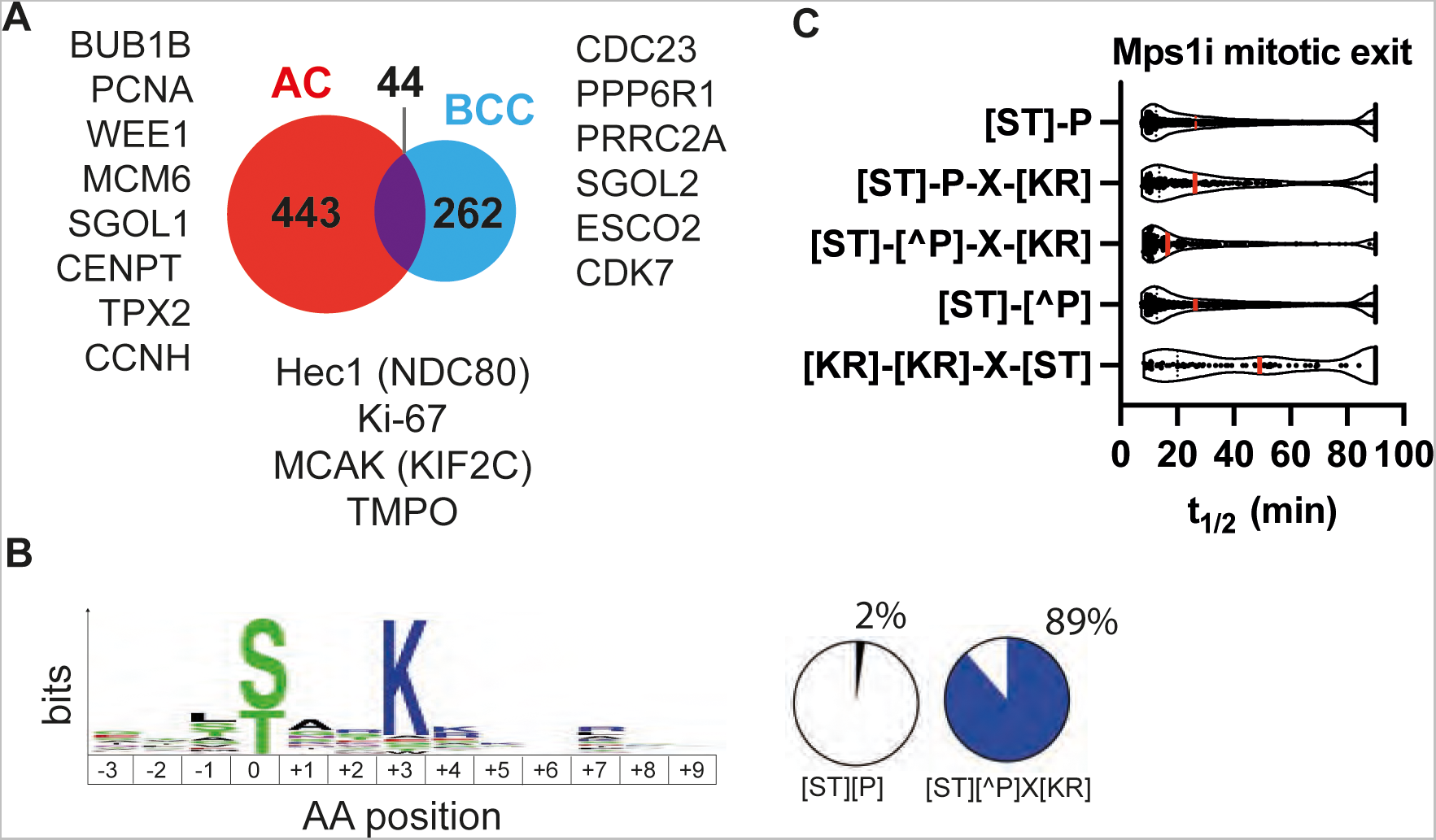
A non-proline directed CDK consensus motif. (A) Overlap between sites reproducibly dependent on Cyclin A (red, N = 2) and Cks1 (blue, N = 2). Phosphorylation is reproducibly enhanced by Cks1, or Cyclin A (compared to Cyclin B-CDK1) for 44 sites (in purple). Selected substrate proteins that are shown for each. (B) Motif enrichment analysis of sites in the overlap (purple shaded area) of (A). (C) Dephosphorylation half-lives for phosphorylation sites detected in Holder et al. 2020 matching the indicated sequence motifs, including proline-directed motifs ([ST]-P, [ST]-P-X-[KR]), the non-proline directed CDK motif identified in (B) ([ST]-[^P]-X-[KR]), sites lacking a +1P ([ST]-[^P]) and sites matching the Aurora consensus ([KR]-[KR]-X-[ST]).

We hypothesized that [ST][^P]X[K] CDK1 sites would be more susceptible to dephosphorylation by protein phosphatases active during prometaphase. This is because PP2A phosphatases show a preference for basic residues C-terminal to the phosphorylated site (Holder et al., 2020), unlike +1P sites, which are generally strongly disfavored by PP2A-B56 (Bancroft et al., 2020; Holder et al., 2020; Kruse et al., 2020).To test this hypothesis, we analyzed a dataset by Holder *et al*. 2020, in which mitotic cells were forced into anaphase by the addition of the Mps1 inhibitor AZ-3146 (Mps1i) and dephosphorylation measured proteome-wide by phosphoproteomics (Holder et al., 2020). The mean half-life (t_1/2_) of [ST][^P]XK sites is ∼17 min, which is significantly shorter than proline-directed sites and non-proline directed sites in general, which both have an average t_1/2_ of ∼28 min (Figure 6C).

We conclude that Cyclin A and Cks1 have overlapping phosphorylation sites characterized by a strong enrichment for a [ST][^P]X[K] motif. These sites are subject to highly dynamic regulation and are rapidly dephosphorylated during forced mitotic exit.

## Discussion

Many cell cycle regulated phosphorylations do not meet the reported consensus sequences for cell cycle kinases, indicating a major gap in our understanding. What are the missing kinases? The results presented suggest a surprising answer: CDK1. In this study, we showed that CDK1 can phosphorylate sites that do not match the classic CDK1 consensus sequence. Furthermore, this non-canonical CDK1 phosphorylation is widespread across the cell cycle regulated phosphoproteome and is regulated by CDK1 subunit composition.

Using an internally controlled, quantitative phosphoproteomics approach, we demonstrated the majority of the mitotic cell cycle regulated phosphoproteome (74%, out of 4,198 sites quantitated) can be directly phosphorylated by CDK1 *in vitro*. To what extent these sites are targets of direct CDK1 phosphorylation is unknown. Even if a fraction of these sites is phosphorylated *in vivo* by other kinases, our data suggest that CDK1 can compensate, or complement the activity of these kinases to phosphorylate these sites in mitosis. Phosphorylation handover from mTOR kinase to CDK1 has been recently described to suppress autophagy in mitosis by targeting the same proline-directed sites in ATG13, ULK1, and ATG14 (Odle et al., 2021). Our data suggest that this handover is likely not limited to mTOR and other proline-directed kinases, and likely extends to non-proline directed kinases. To what extent this handover occurs, and how this is regulated is an open question that will be important to address.

Differential substrate targeting is regulated by subcellular localization of Cyclin A and Cyclin B in living cells (Jackman et al., 2002; Moore et al., 2002). However, in fixed cells, nucleocytoplasmic transport is inactive and in principle, these complexes have equal access to cellular substrates. Compared to Cyclin B-CDK1, substrate proteins preferentially phosphorylated by Cyclin A-CDK1 differ in annotated functions and subcellular localization, consistent with the observations that substrate targeting is conferred by the Cyclin subunit. We observed no significant difference in RXL motifs between Cyclin A- and Cyclin B-dependent substrates, suggesting that there are sequence elements within substrate proteins encoding for specificity. These additional sequence elements, like the RXL motif, could be encoded in *cis* (i.e., within the same protein sequence as the phosphoacceptor residue). Because subcellular organization and protein-protein interactions are largely retained in fixed cells, it is possible that substrate choice in these assays is determined by interactions in *trans*. For example, high affinity or avidity interactions between CDK1 complexes with a scaffolding protein could promote phosphorylation of other substrates proximal in space. Spatial determinants of substrate phosphorylation are key, for example, in models for the regulation of kinetochore-microtubule attachments by kinases like Aurora B and Plk1 (Samejima et al., 2015; Singh et al., 2021).

We show that substrate specificity *in vitro* is altered by the Cyclin subunit and by CDK1 concentration. Both Cyclin A-CDK1 and Cyclin B-CDK1 show similar concentration-dependent changes in substrate specificity (Figure 2C, Supplementary Figure 1F). These data suggest sites with highest sensitivity towards Cyclin B-CDK1 are likely to be also phosphorylated by Cyclin A-CDK1. However, at higher Cyclin-CDK1 concentrations, Cyclin A-CDK1 preferentially phosphorylates specific sites to a much higher extent (i.e., >6.5-fold, Figure 1E), as compared with Cyclin B-CDK1, and vice versa. Taken together, these data suggest that in cells, the differential requirement for Cyclin B versus Cyclin A might only arise for phosphorylations that require high CDK1 activity. And as a corollary, Cyclin B and Cyclin A might be functionally redundant for substrate phosphorylation with low CDK1 activity thresholds.

The *in vitro* experiments show an enrichment for threonine phosphoacceptor residues for CDK1 (Figure 1C) and we observe a positive correlation between CDK1 sensitivity and phosphothreonine frequency (Figure 2F). The yeast Cdk1 homologue, cdc28, shows a slight preference for serine over threonine phosphoacceptor residues *in vitro* in peptide assays (Chen et al., 2014). Human CDK1 is anticipated to have the same preference for serine based on the conserved DFG+1 residue (Leucine), making phosphoacceptor preference by the kinase an unlikely explanation for threonine enrichment (Chen et al., 2014).

An alternative explanation could be differential phosphorylation occupancy of serine and threonine residues in fixed G2&M cells. The PP2A:B55 phosphatase has been shown to prefer dephosphorylating phosphothreonine residues with a +1 Proline (Cundell et al., 2016). PP2A:B55 is active in interphase, inactivated at mitotic entry, and reactivated at mitotic exit. Consistent with this idea, CDK1 sensitivity is negatively correlated with endogenous phosphorylation (Figure 2G) and kinase assays on phosphatase treated cells show a significant reduction in phosphothreonine CDK1 sites (Supplementary Figure 3E). Clusters 1 and 2, which are likely to be most enriched in PP2A:B55 target sites, are not differentially enriched in proteins with cell cycle functions. This result is consistent with the idea that PP2A:B55 broadly antagonizes CDK1 activity in interphase (Krasinska et al., 2011). Interestingly, phosphorylation sites in these two clusters are the most sensitive to CDK1 phosphorylation *in vitro* (Figure 2A) and *in vivo* (Figure 2I), and likely targeted for extensive dephosphorylation by protein phosphatases (Figures 2F, H). These results suggest that despite being most sensitive to CDK1 phosphorylation, these sites are not phosphorylated at appreciable levels due to active interphase phosphatases. The data therefore support a model whereby phosphatase inactivation being a key driver of CDK1 substrate phosphorylation.

Using G2&M cells with endogenous phosphorylation for fixed cell assays mimics the context *in vivo*, where a subset of phosphothreonine-biased CDK1 sites will be targeted for dephosphorylation by PP2A:B55 and be exquisitely sensitive to phosphorylation when CDK1 activity rises and PP2A:B55 is inactivated at the G2 to M transition. In future, however, it will be important to measure CDK1 phosphorylation sensitivity in phosphatase pre-treated fixed G2&M cells, which will fully address if quantitative changes in CDK1 is sufficient to enforce phosphorylation order.

The fixed cell assays we have developed to understand CDK1 regulation can be extended to other cellular enzymes that produce a protein mass modification measurable by mass spectrometry, e.g., ubiquitination. In contrast to *in vitro* assays on cell lysates, it is straightforward to perform sequential enzymatic reactions on fixed cells. We have capitalized on this feature of our assays to study the role of priming phosphorylation on CDK1 phosphorylation proteome-wide (Figure 4). Sequential reactions can be used to study crosstalk between protein post-translational modifications in a highly controlled manner that is challenging in living cells with active negative and positive feedback mechanisms.

Several proteins with the [ST][^P]XK motif have known roles in mitotic regulation, including Ki-67, BubR1 and MCAK. Ki-67 is localized to mitotic chromosomes and is an important scaffolding factor to form the mitotic chromosome periphery (Booth and Earnshaw, 2017; Cuylen-Haering et al., 2020). Ki-67 functions as a biomolecular surfactant, facilitating chromosome dynamics during mitosis and promoting timely chromosome segregation (Cuylen et al., 2016). 24 CDK1 phosphorylation sites on Ki-67 are Cks1-dependent (out of 49 CDK1 sites detected in total). Many phosphorylation sites on Ki-67 are Cyclin B-dependent in cells (Hégarat et al., 2020). Acute depletion of Cyclin B causes loss of Ki-67 at the chromosome periphery and defects in chromosome segregation (Hégarat et al., 2020). It will be interesting to test if these effects are Cks1-dependent, as would be predicted from our data.

Non-proline directed CDK1 phosphorylation has been previously reported (Blethrow et al., 2008; Michowski et al., 2020). Recently, it was shown that ∼30% of direct CDK1 phosphorylation sites were non-proline directed in mESCs (Michowski et al., 2020). Interestingly, these sites showed an enrichment for a C-terminal basic residue, consistent with previous reports on individual substrates and the [ST][^P]XK motif described in our study (Suzuki et al., 2015). We speculate these sites are especially important for spatial and temporal regulation in mitosis because they likely require high avidity interactions for phosphorylation and are exquisitely sensitive to dephosphorylation (Figure 6C), thereby providing a wide dynamic range for rapidly tuning protein function.

In this study, we show that the qualitative nature of the CDK1 complex, namely the subunit composition, has a striking effect on the phosphorylation consensus sequence. The regulated consensus sequence switch shown in this study highlights the importance of site-level phosphorylation analysis enabled by mass spectrometry-based phosphoproteomics. CDK1/2 substrates are phosphorylated in both interphase and in mitotic cells, but on distinct sites that have differential impact on protein function. Proteins in the Mcm family (Mcm1-7), which form the replicative helicase on chromatin to support DNA replication, are excellent examples of this. *Xenopus laevis* Mcm4 (*xl*Mcm4) is hyperphosphorylated in mitosis when replicative helicases are inactive. Complete dephosphorylation of *xl*Mcm4 prevents chromatin binding, whereas pre-replicative complexes bound to chromatin are hypo-phosphorylated, i.e., showing an intermediate level of phosphorylation between dephosphorylated and hyperphosphorylated (Pereverzeva et al., 2000). In our analysis, two Mcm4 sites are detected, T23 and S120, which show high and low sensitivity to CDK1 phosphorylation *in vitro*. Interestingly, S120 meets the S[^P]XK consensus sequence for non-proline directed CDK1 phosphorylation, and is highly phosphorylated in mitosis (∼84% stoichiometry) (Olsen et al., 2010).

Our study provides evidence that these non-proline directed sites are not due to adventitious binding and instead are likely to have specific functions in cells on the basis of their differential regulation by CDK1 subunit composition (Figures 1 and 3) and by phosphatases (Figure 6C). Non-proline directed phosphorylations might be a consequence of long substrate residence times due to high avidity docking interactions between the non-catalytic subunits (Cyclin, Cks1) and substrate. In future, it will be important to address the structural and molecular basis for phosphorylation consensus switching and to assess if this consensus switching is a general mechanism for kinase regulation.

## Acknowledgements

We thank members of the Ly, JP Arulanandam, Julie Welburn, Constance Alabert, Julian Blow and Bill Earnshaw groups for helpful discussions. We thank the FingerPrints Proteomics Facility and Edinburgh Protein Production Facility for technical support. This work is supported by a Wellcome Trust and Royal Society Sir Henry Dale Fellowship to T.L. (206211/Z/17/Z), a Darwin Trust PhD studentship to A. A., a UK Medical Research Council Programme Grant to J.E. (MR/N009738/1), core funding for the Wellcome Centre for Cell Biology (091020), a Wellcome Multi-User Equipment Grant to T.L. (218305/Z/19/Z) and a Wellcome Innovation Award for mass spectrometry equipment (218448/Z/19/Z).

## Author contributions

TL: Conceptualization, Supervision, Formal Analysis, Writing – original draft, Funding acquisition, Investigation, Writing – original draft, Writing – review & editing

AA: Conceptualization, Investigation, Formal Analysis, Methodology, Writing – original draft, Writing – review & editing

SK: methodology (protein expression and purification), investigation and writing (original draft)

JE: Conceptualisation, resources, supervision, funding acquisition, writing original draft, review and editing.

## Supplementary Figure Legends

**Supplementary Figure 1.**
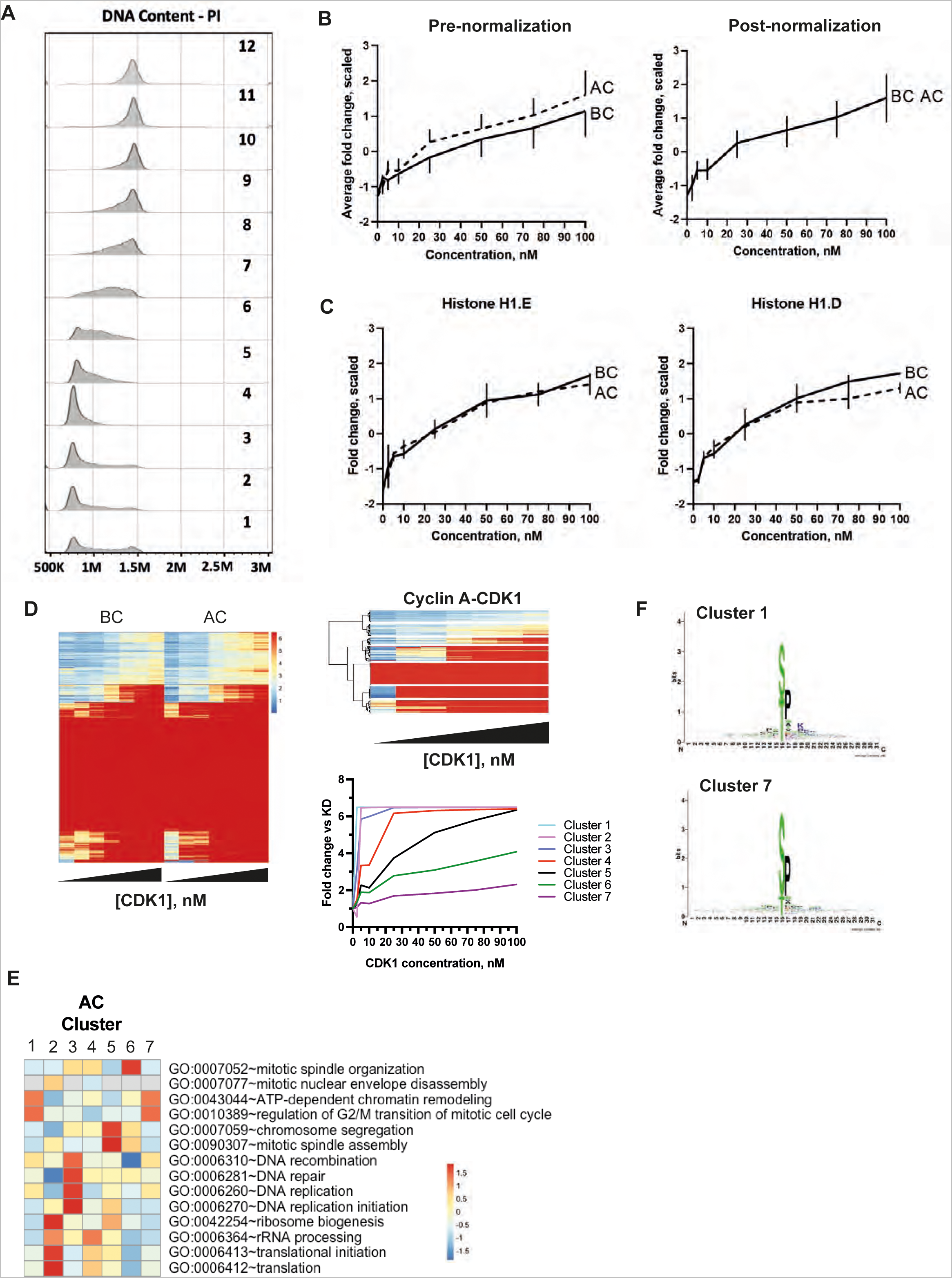
(A) Flow cytometry data showing enrichment of cell cycle phases in fractions collected by centrifugal elutriation. (B) Pre- (left) and post- (right) normalization scaled fold change comparing AC and BC for all CDK1 phosphorylation sites. (C) Post-normalization scaled fold changes for histone H1, AC vs BC. (D) Hierarchal clustering identified seven clusters that have decreasing sensitivity towards CDK1 phosphorylation (1 and 7 being most and least sensitive, respectively). (E) Consensus motif for clusters with most and least sensitive Cyclin A-CDK1 sites. (F) Heatmap showing selected enriched functional annotations for each cluster in (D). Colour indicates scaled fold enrichment.

**Supplementary Figure 2.**
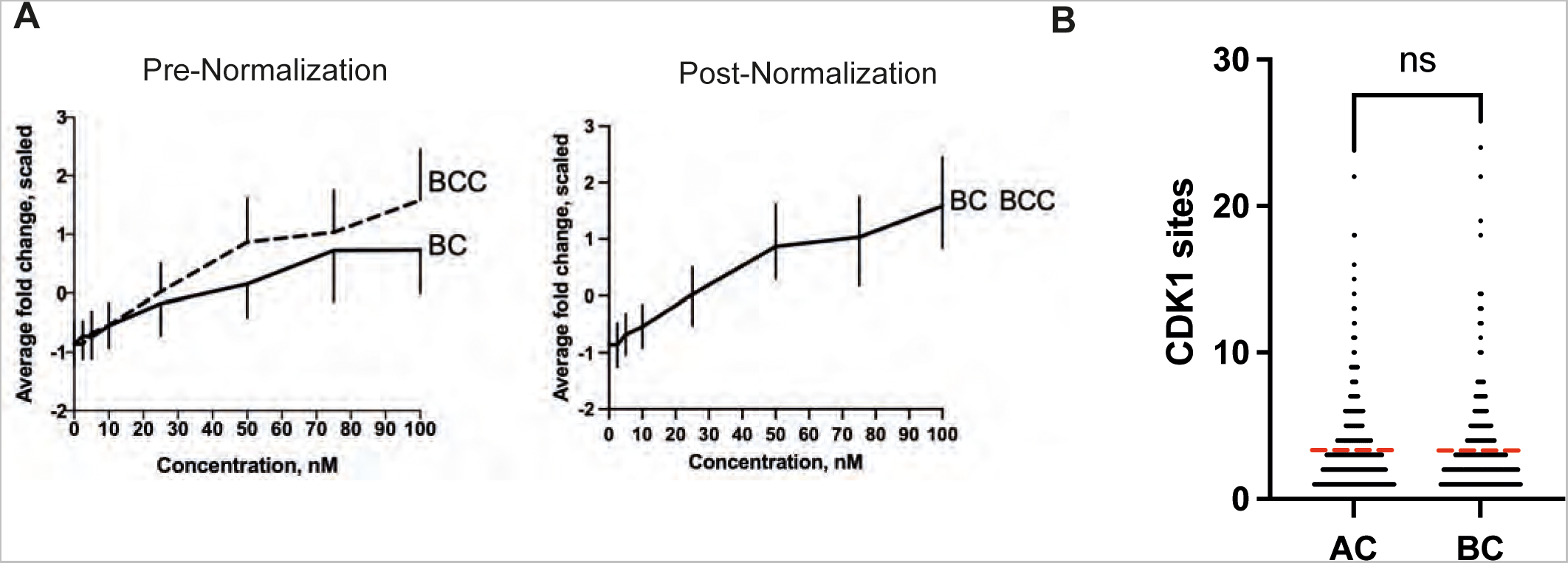
(A) Pre- (left) and post- (right) normalization scaled fold change comparing BC and BCC for all CDK1 phosphorylation sites. (B) Comparison of multisite phosphorylation between Cyclin A- and Cyclin B-dependent substrates.

**Supplementary Figure 3.**
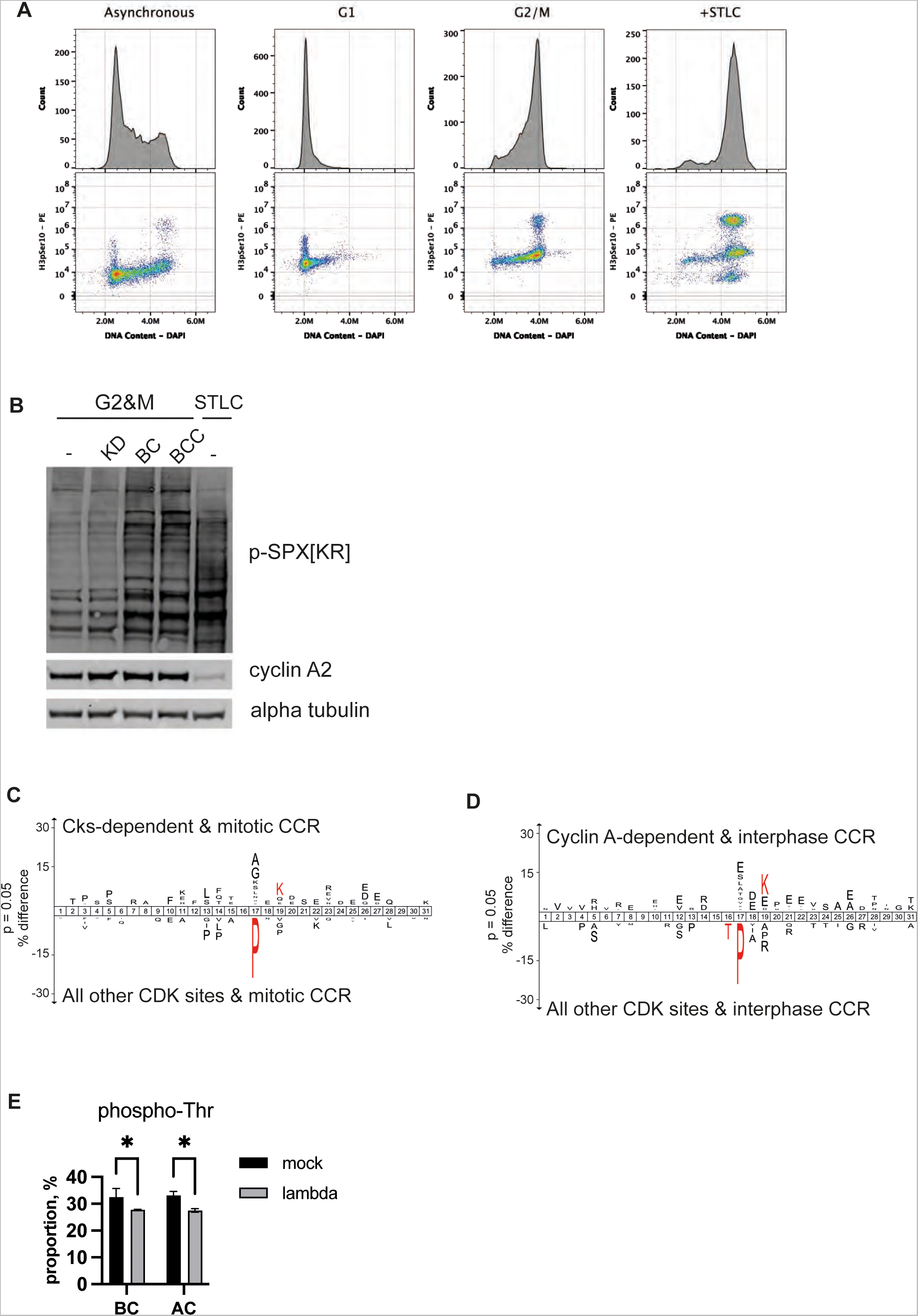
(A) Flow cytometry data showing enrichment of cell cycle phases in fractions collected by centrifugal elutriation and STLC mitotic arrest. (B) Immunoblot analysis of G2&M-phase cells phosphorylated by indicated kinase complexes, in comparison with STLC-arrested mitotic cells. (C) Motif enrichment analysis of phosphorylation sites that are Cks1-dependent *in vitro* and mitotic CCR. (D) Motif enrichment analysis of phosphorylation sites that are Cyclin A-dependent *in vitro* and interphase CCR. (E) Proportion of phosphothreonine CDK1 sites comparing mock- and lambda phosphatase-treated G2&M cells phosphorylated with the indicated kinases.

## Supplementary Tables

Supplementary table 1: Comparison of AC and BC

Supplementary table 2: Titration of CDK1 activity

Supplementary table 3: Comparison of BC and BCC

Supplementary table 4: Priming phosphorylation for Cks1

Supplementary table 5: BCC specific Mitosis regulated sites

Supplementary table 6: AC specific interphase regulated sites

Supplementary table 7: Overlap between BCC and AC substrates

## STAR Methods

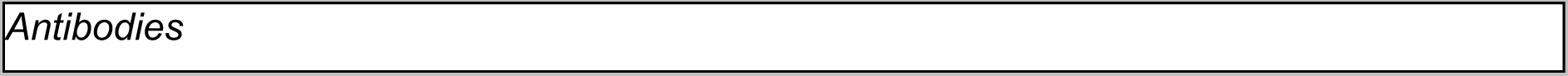
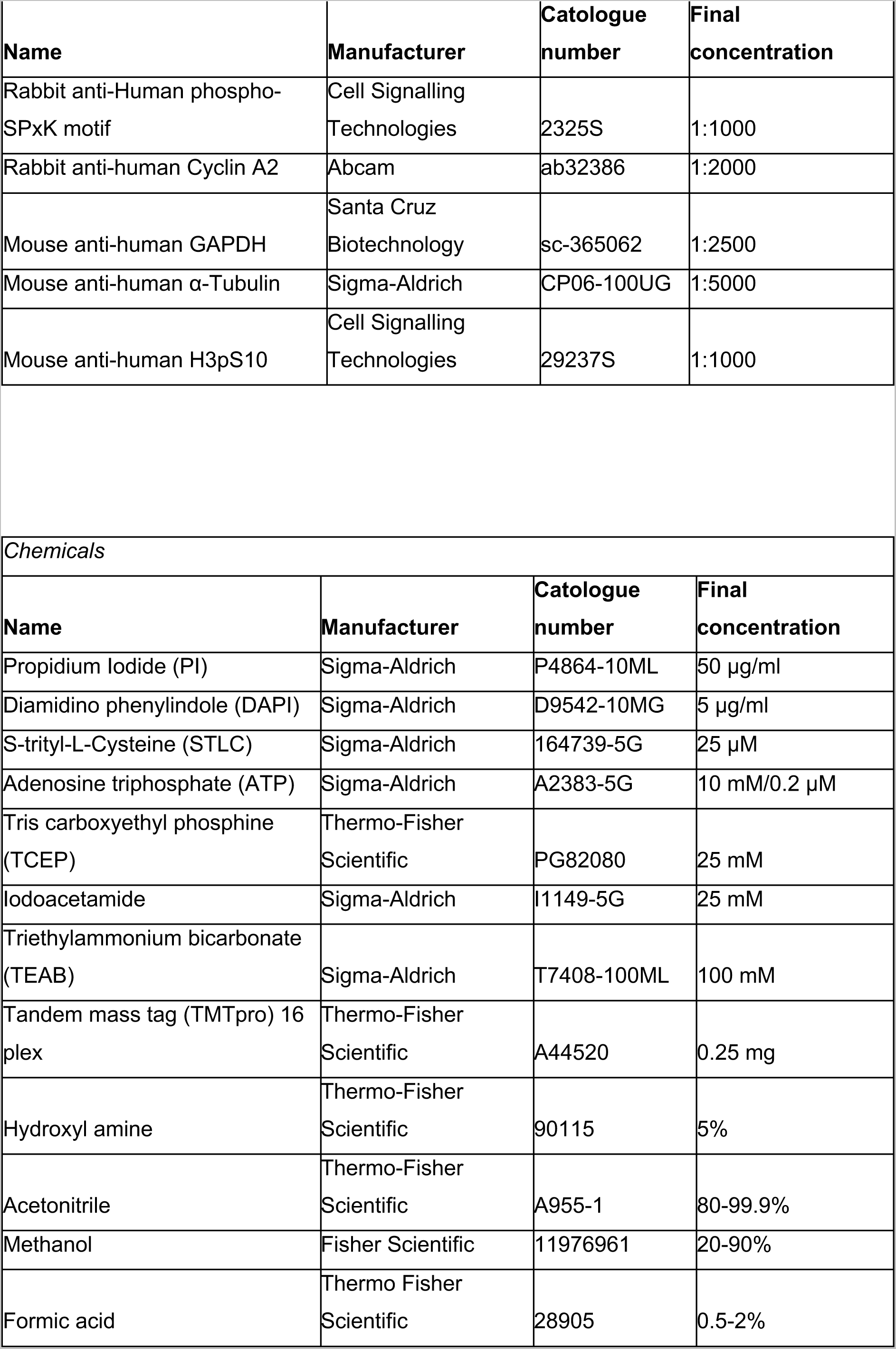
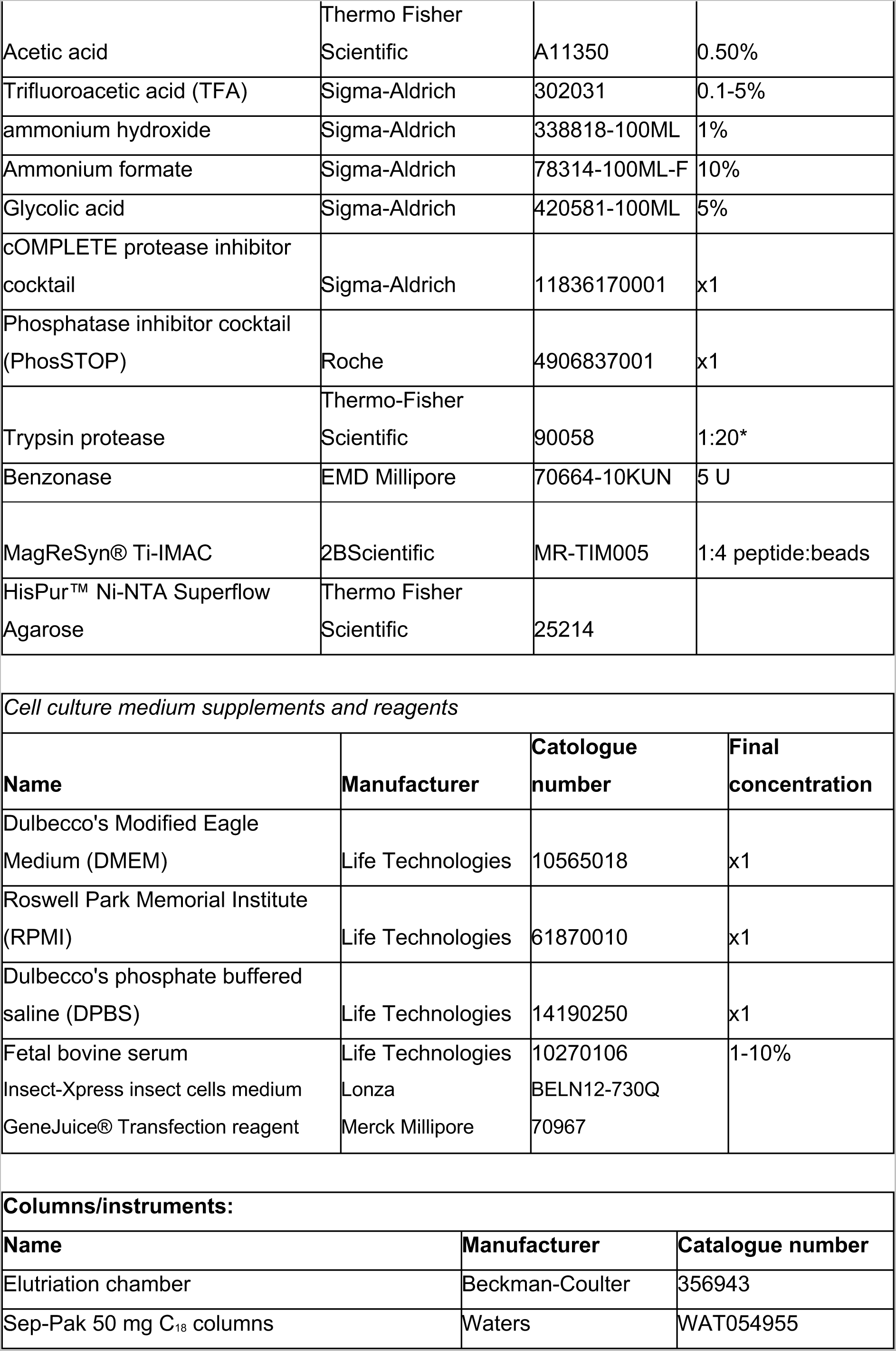
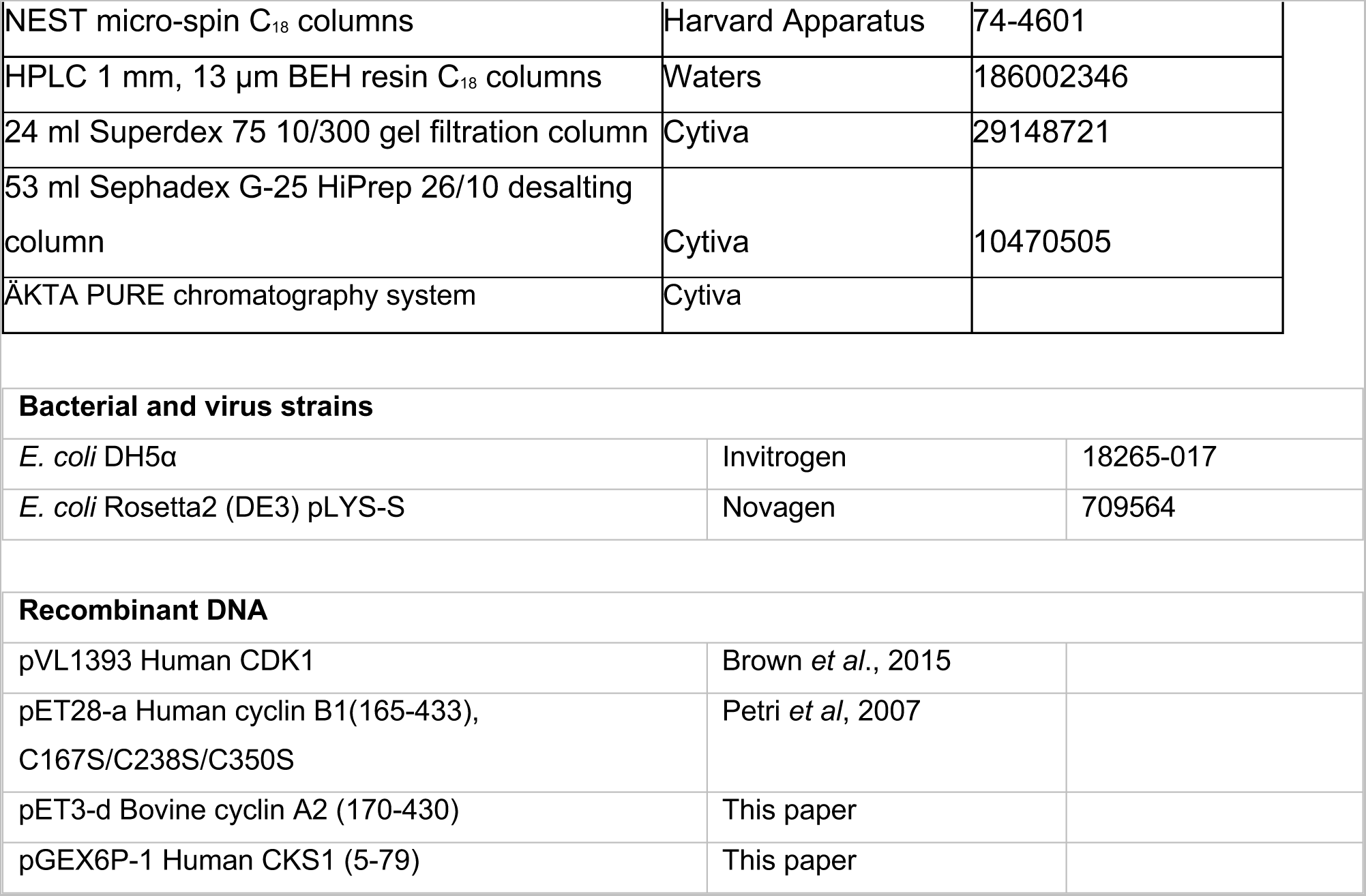

### Cell culture and centrifugal elutriation

TK6 cells were seeded at 80,000 cells/ml in 50 ml of Roswell Memorial Park Institute (RPMI) tissue culture medium supplemented with 10% Fetal Bovine Serum (FBS) in eight 15 cm dishes. After 48 hours, cells were centrifuged, pooled, washed once in Dulbecco’s Phosphate Buffered Saline (DPBS), and resuspended in 1% Formaldehyde. Cells were mixed on a rotator at room temperature for 10 mins. Cells were washed in DPBS and permeabilized in 90% Methanol for a minimum of 24 hours at -20 °C. For centrifugal elutriation, methanol was removed, and cells were resuspended in elutriation buffer (5 mM MES, 100 mM NaCl, 1% FBS) and placed in elutriation chamber fitted into ultra-centrifuge. Cells were trapped in the elutriation chamber at 2010 rpm and 15 ml/min flow driven by a peristaltic pump. Cell size fractions were collected by progressively increasing the flow rate up to 35 ml/min. The cell cycle phase distribution of each elutriation fraction was analyzed by staining cells with propidium iodide (50 µg/ml RNAse A, 50 µg/ml Propidium Iodide in DPBS) for 30 mins prior to flow cytometry analysis. Fractions were combined in DPBS supplemented with 1x Roche phosphatase inhibitor cocktail to obtain pooled fractions enriched in either 2N or 4N DNA content. A sample of cells was immunostained with H3S10ph antibody conjugated to Phycoerythrin (PE) for 30 mins. Cells were washed once and resuspended in flow buffer containing 5 µg/ml DAPI. To arrest cells in mitosis, TK6 cells were seeded at 800,000 cell/ml in RPMI supplemented with 10% FBS and 25 µM STLC. After 16 hours, cells were then harvested, fixed and permeabilized as described above.

### Protein expression and purification

Full-length human CDK1 was expressed in insect cells from pVL1393 as a 3C-protease cleavable GST fusion, which leaves a short cloning artefact (GPLGS) at the N terminus. This construct was expressed, purified and phosphorylated (using GST-CAK1) as previously described(Brown et al., 2015; Brown et al., 1999b). T161 phosphorylated CDK1 was then purified from GST-CAK1 by size exclusion column chromatography on a Superdex 75 26/60 column equilibrated in 50 mM Tris pH7.5, 150 mM NaCl, 0.5 mM TCEP. Human cyclin B1, residues 165–433 carrying the C167S/C238S/C350S mutations, was expressed in recombinant *E. coli* cells and purified as described exploiting the thrombin-cleavable hexa-histidine tag encoded by the pET28-a (+) vector (Petri et al., 2007). Human Cks1 was expressed from pET21a in *E. coli* cells and purified as described (Brown et al., 2015).

Bovine cyclin A2, residues V170-V430 was expressed in *E. coli* Rosetta2 (DE3) pLYS-S cells as a GST-fusion from a modified pET3-d vector. It was purified by affinity purification followed by 3C cleavage to remove the GST tag and then a subsequent size-exclusion chromatography step (Superdex 200 16/60 column). The bovine cyclin A2 has the GPLMKY sequence at the N-terminus as a cloning artefact following 3C cleavage. As previously described (Brown et al., 1995), bovine cyclin A2 was purified in buffer containing MgCl_2_ (300 mM NaCl, 100 mM MgCl_2_, 50 mM Tris-HCl, pH 8.0, 1 mM DTT) to help prevent aggregation.

To prepare the binary T161pCDK1-cyclin B1 and ternary T161pCDK1– cyclin B1-CKS1 complexes, components were individually purified and then mixed in molar excesses of cyclin B1 and CKS1 over CDK1 as required, and essentially as described (Brown et al., 2015). The interaction between CDK1 and cyclin B1 is dependent on the concentration of salt in the buffer. In each case, the final step to assemble the complex was carried out on a Superdex 75 HR26/60 SEC column equilibrated in modified Tris-buffered saline containing 1.0 M NaCl, 50 mM Tris-HCl, pH 8.0, 1 mM DTT. T161pCDK1-cyclin A2 was prepared by mixing purified phosphorylated CDK1 with an excess of purified bovine cyclin A2 and separating the complex by size exclusion chromatography on a Superdex 75 HR26/60 column equilibrated in 50mM Tris pH 8.0, 200mM NaCl, 1mM DTT. For each purification, fractions containing the desired complex were pooled and concentrated to circa 10–12 mg ml^-1^ by ultrafiltration and then fast frozen in aliquots in liquid nitrogen before storage at -80 °C.

A recombinant complex comprised of a truncated Aurora B (55-344) and the C-terminus of INCENP (835-903) was expressed and purified from bacterial cells. To co-express these proteins, Escherichia coli Rosetta cells were co-transformed with plasmids carrying the open reading frame sequences of both the truncated INCENP and the truncated, N-terminally 6x Histidine-SUMO tagged, Aurora B and grown on LB agar plate (Supplemented with Kanamycin, Chloramphenicol and Spectinomycin antibiotics). Cultures were scaled up and grown at 37°C by inoculating 20 ml of cells from a dense bacterial suspension into 2 Litres of LB growth medium supplemented with the antibiotics described above. Once bacterial cells reached a density of approximately 0.5 OD, the temperature was decreased to 18°C, and the expression was induced by adding 350 µM IPTG for 18 hours. Cells were then pelleted and lysed with a bio-disruptor in lysis buffer (25 mM HEPES, 500 mM NaCl, 25 mM Imidazole, 2 mM β-mercaptethanol, 1 x cOMPLETE protease inhibitor cocktail and 50 U Benzonase; pH 7.5). To pull down the expressed Aurora B complex, supernatant containing the soluble proteins was collected by spinning lysates at 22,500 rpm for 50 mins at 4°C and incubated with Nickel coated silica beads for 2 hours at 4°C. To remove non-specifically bounds proteins, washes with lysis buffer, chaperone buffer (25 mM HEPES, 1000 mM NaCl, 30 mM Imidazole, 50 mM KCl, 10 mM MgCl_2_, 2 mM ATP and 2 mM β-mercaptethanol; pH 7.5) then a low salt buffer (25 mM HEPES, 200 mM NaCl, 25 mM Imidazole and 2 mM β-mercaptethanol; pH 7.5) were carried out by resuspending beads in each and centrifugation at 500 g for 5 mins at 4°C. To elute the Aurora B complex, beads were dipped in Imidazole elution buffer (25 mM HEPES, 200 mM NaCl, 500 mM imidazole, 2 mM β -mercaptethanol) for 2 hours at 4°C then centrifuged as described above. This was followed by four rounds of resuspension then centrifugation in the Imidazole elution buffer and pooling of the eluent. To cleave the 6x Histidine-SUMO tag from the recombinant Aurora B complex, samples were first passed through a 50 ml pre-packed desalting column fitted on ÄKTA liquid chromatography instrument and fractionated in a dialysis buffer (25 mM HEPES, 200 mM NaCl and 2 mM β-mercaptethanol; pH 7.5) to remove the Imidazole. Desalted proteins were then incubated overnight at 4°C with the protease SENP2 to cleave the tag. To purify the Aurora B complex, samples were concentrated to 300 µl and loaded into a pre-packed, 24 ml Superdex 75 10/300 gel filtration column fitted on ÄKTA instrument in gel filtration buffer (25 mM HEPES, 200 mM NaCl, 4 mM Dithiothreitol [DTT] and 5% Glycerol; pH 7.5).

Fractions containing the two subunits of the recombinant complex were identified by SDS-PAGE, snap frozen in liquid nitrogen and stored at -80°C until the *in vitro* kinase assays.

### In vitro kinase assays on fixed cells

To induce protein phosphorylation *in vitro*, 2 million fixed and permeabilized TK6 cells from the pooled elutriation fractions above were blocked in 40 mM Tris (supplemented with 5% BSA and 1x Roche phosphatase inhibitor cocktail) for 10 mins. Cells were then washed by centrifugation at 15,000 g for 30 secs and placed in 400 µl of phosphorylation master-mix (40 mM Tris, 0.5% BSA, 10 mM ATP, 1x Roche phosphatase inhibitor cocktail and recombinant CDK1 complexes) for 40 mins at room temperature. For phosphorylation with Aurora B, cells were blocked in DPBS (Supplemented with 5% BSA and 1x Roche phosphatase inhibitor cocktail) for 10 mins on ice then placed in a master-mix (DPBS, 0.5% BSA, 10 mM ATP, 1x Roche phosphatase inhibitor cocktail and recombinant Aurora B) for 40 mins at 37°C. To stop the phosphorylation reactions, cells were quenched by washing three times with 11 mM EDTA in either 40 mM Tris (Supplemented with 0.5% BSA) for assays with CDK1 or in DPBS (Supplemented with 0.5% BSA) for assays with Aurora B. Cells were then resuspended in 300 µl of ice cold DPBS containing 1x Roche phosphatase inhibitor cocktail. To check the *in vitro* phosphorylation by Western blotting, 200 µl of this suspension was placed in a new tube, centrifuged and pellets were resuspended in 70 µl cell extraction buffer (1 mM HEPES, 10 µM EDTA, 2% SDS, 1x cOMPLETE protease inhibitor cocktail and 1x Roche phosphatase inhibitor cocktail). Extracts were then sonicated for 30 secs at 10% amplitude, and crosslinking was reversed by heating at 95°C for 50 mins.

Proteins in the lysates were then reduced by adding 25 mM TCEP and mixed with 25 µl of LDS sample buffer for loading. Proteins were separated by SDS-PAGE at 150 V in 1x NuPAGE MES Running Buffer for 2 hours and transferred onto 0.2 µm nitrocellulose membranes at 0.2 A in 80% NuPAGE MES buffer + 20% methanol (v/v) for 2 hours. Membranes were stained overnight with anti-phospho-SPxK motif and anti-Tubulin antibodies in Tris buffered saline (TBS, Supplemented with 5% BSA) at 4°C. This was done after blocking with TBS (Supplemented with 5% milk) for a minimum of 1 hour. To remove the primary antibodies, membranes were washed three times with TBS-T (TBS and 0.1% Tween) for 5 mins each. Secondary antibodies conjugated to IRDye680 or IRDye800 in TBS (Supplemented with 5% BSA) were then added for 1-2 hours at room temperature and bands were visualized by scanning the membranes with Li-COR Odyssey instrument.

### Sample preparation for TMT phosphoproteomic analysis of fixed cells

To prepare peptide digests from fixed cells, 100 µl of cells from above were washed by centrifugation, resuspended in digestion buffer, which consisted of: 100 mM triethyl ammonium bicarbonate (TEAB), 2 mM MgCl_2_ and 5 U Benzonase; pH 8.5. Nucleic acids were digested at 37°C for 30 mins (Kelly et al., 2022). Proteins were then digested by adding 1.25 µg Trypsin protease for 16 hours at 37°C followed by another addition of 1.25 µg trypsin for 4 hours. Peptides were then acidified by adding formic acid to a final concentration of 2% and desalted using NEST C_18_ micro-spin columns. Briefly, peptides were bound to C_18_ columns that were previously conditioned with 100% acetonitrile and equilibrated with 0.5% formic acid. Columns were then washed twice with 0.5% formic acid and peptides were eluted with 80% acetonitrile diluted in 0.5% formic acid. To remove solvent, samples were dried at 30°C until fully dry. Peptides were resuspended in 50 µl of 100 mM TEAB and mixed with 0.25 mg of TMTpro from a set of 16 plex resuspended in 10 µl acetonitrile for 1 hour. The isobaric labelling reaction was then quenched by adding 2.5 µl of 5% hydroxylamine to these samples for 15 mins at 37°C. Peptides from samples in the following experiments were pooled together: AC/BC titration experiment discussed in Figures 1 and 2; BCC and BC titration experiment in Figure 3. The experiment involved λ Phosphatase pre-treatment of fixed cells in Figure 4 was pooled in the same TMT set with samples for CCR sites identification discussed in Figure 5 and the identification of mitotic CCR sites that were Cks1 dependent was done with the same samples used for identifying the priming phosphorylation kinase in Figure 4. Peptides from each pool were then dried, resuspended in 0.5% formic acid, and divided into two fractions each was desalted in a 50 mg Sep-Pak C_18_ column. To remove free TMT from samples, an extra wash with 0.5% acetic acid was added to the protocol and elution was done in 80% acetonitrile, this time diluted in 0.5% acetic acid. Samples were phosphoenriched by mixing peptides with 3.2 mg of MagReSyn® Ti-IMAC in load buffer, which consisted of 80% acetonitrile, 5% trifluoroacetic acid (TFA) and 5% glycolic acid, for 20 mins at 25°C. Beads were washed for 2 min with 80% acetonitrile + 1% TFA. This was followed by two 2 min washes in 10% acetonitrile + 0.2% TFA. Phosphorylated peptides were eluted in 1% ammonium hydroxide for 15 mins twice in elution buffer. This was followed by a second elution for 1 hour in a 50/50 mix of 1% ammonium hydroxide and acetonitrile (v/v). To increase the number of phosphorylated peptides identified, the flow through was dipped in a new batch of MagReSyn® Ti-IMAC beads and the phospho-enrichment step was repeated as described above. 5% of the pooled peptides were kept without phospho-enrichment for total proteome analysis. To remove any residual magnetic beads, peptides were dried and desalting with NEST micro-spin C_18_ columns was performed as described above. For deep phosphorylation analysis, peptides were fractionated using high pH reverse phase HPLC. Briefly, peptides were passed through a 1 mm column packed with 13 µm sized BEH silica resin coated with C_18_ and were eluted with a gradient of 15 – 80% B, with the following A and B mobile phases: 10 mM ammonium formate pH 9.3, 10/90 mixture of 10 mM ammonium formate pH 9.3 and 100% acetonitrile. Peptides were eluted into 16 wells, dried and stored at -20°C until data acquisition by mass spectrometry.

### Proteomics data analysis

In the kinase assay where phosphorylation by BC was compared to BCC described in Figure 1, the TMTpro labelled and phosphoenriched samples were analyzed using Dionex Ultimate 3000 HPLC-Coupled Tribrid Fusion Lumos mass spectrometer. Samples were loaded and separated using 75 μm × 50 cm EASY-Spray column with 2 μm sized particles, which was assembled on an EASY-Spray source and operated constantly at 50°C. Two mobile phases were used to separate the peptides: Phase A consisting of 0.1% formic acid in LC-MS grade water and phase B consisting of 80% acetonitrile and 0.1% formic acid. Peptides were loaded onto the column at a flow rate of 0.3 μL/min and eluted at a flow rate of 0.25 μL/min according to the following gradient: 2 to 40% mobile phase B in 120 min and then to 95% in 11 min. Mobile phase B was retained at 95% for 5 min and returned back to 2% a minute after until the end of the run (160 min in total for each fraction). A voltage of 2.2 kV was set when spraying this gradient of peptides into the front end of the mass spectrometer at ion capillary temperature of 280°C with a maximum cycle time of 3 secs. An MS1 scan at a resolution of 120,000 in Orbitrap detector was performed with a maximum injection time of 50 msec and the top 10 most abundant ions within a scan range of 380-1500 m/z based on the m/z signal with charge states of 2-6 from that scan were chosen for fragmentation in a HCD cell at 28%. This was followed by a rapid MS2 scan on a linear ion trap for peptide identification with a maximum injection time of 50 msec. To minimize the TMTpro reporter ion ratio distortion, 5 precursor fragments were selected for a synchronous precursor selection (SPS) MS3 method from 3 precursor dependent scans (McAlister et al., 2014). These fragments were further fragmented at 55% collision energy in a HCD chamber and analyzed using an Orbitrap detector at a resolution of 55,000 with a 90 msec maximum injection time. The samples from this experiment that were not phosphoenriched were analyzed for total proteome analysis using the same method, except that the MS2 fragmentation was performed in a CID cell at 35% energy.

In the experiment where AC and BC phosphorylation of fixed cells was compared (Figure 3), both total and phosphorylated peptides were loaded into a trap column (100 μm × 2 cm, PepMap nanoViper C18 column, 5 μm, 100 Å) attached to a Dionex Ultimate 3000 RS system in 0.1% TFA for 3 mins and then separated through analytical column (75 μm × 50 cm, PepMap RSLC C18 column, 2 μm, 100 Å) in the following mobile phases: 0.1% formic acid (Solvent A) and 80% acetonitrile + 0.1% formic acid (Solvent B). Separation was carried out using a linear solvent gradient of 5% to 35% for 130 mins followed by a steep gradient to 98% up until 152 mins after which the solvent concentration was dropped back to 5%. The separation was carried out at a flow rate of 300 nl/min. Peptides were then sprayed into the front end of a Tribrid Fusion mass spectrometer through a nanoelectrospray ionizer with a cycle time of 3 secs and the MS1 data for precursor ions were acquired in an Orbitrap detector at a resolution of 120,000. The top 10 most abundant peaks with charge states of 2-6 were then fragmented in a HCD chamber at 28% and analyzed on a linear ion trap for peptides identification in a maximum injection time of 70 msec. A neutral loss that matches the molecular weight of the phosphate group (98 m/z) was set to identify phosphorylated peptides. SPS MS3 was then performed on the top 5 precursor fragments from 5 precursor dependent scans following HCD fragmentation (58%) in Orbitrap detector at a resolution of 50,000 with a maximum injection time of 110 msec.

Finally, phosphorylated and total samples that involved sequential reactions with phosphatase and CDK1 described in Figure 2 and those used for the analysis of CDK1 CCR sites described in Figure 4 were labelled with TMTpro and pooled into the same set and analyzed on an Orbitrap Eclipse mass spectrometer. Peptides were initially trapped in PepMap nanoViper C18 column (100 μm × 2 cm, 5 μm, 100 Å) in 0.1% TFA for 5 mins then fractionated with analytical PepMap RSLC C18 column (75 μm × 50 cm, 2 μm, 100 Å) on a Dionex Ultimate 3000 RS system with a 5%-35% gradient for 130 mins. This was followed by a steep increase in solvent concentration to 98% for up to 152 mins then a drop to 5% for 1 min. peptides from the gradient were injected into the front end of a Tribrid Eclipse mass spectrometer through a nanoelctrospray ionizer in a 3 secs cycle time. Precurosr ions were detected in a master scan using Orbitrap detector at a resolution of 120,000 with a maximum injection time of 50 msecs. Precursor ions with top 10 signals and charge states of 2-7 were selected for fragmentation using HCD (28%) and analyzed by a linear ion trap with a maximum injection time of 50 msecs. SPS-MS3 of 5 fragments from 5 precursor dependent runs were fragmented by HCD (55%) and analyzed by Orbitrap at a resolution of 50,000 with a maximum injection time of 90 msecs.

Raw data from the assays on fixed cells were processed on MaxQuant version 1.6.14 (Cox and Mann, 2008). To normalize the data, reporter intensities for histone proteins detected in all TMTpro channels in the Protein Groups file were summed. Summed intensities were used to normalize the reporter ion intensities for sites in the corresponding TMTpro channels in the PhosphoSTY file to adjust for mixing error. The fold change in phosphorylation intensity for each site was then calculated by dividing the normalized reporter ion intensity of that site in the sample treated with the active recombinant kinase by its normalized reporter ion intensity in the control sample. Sites with at least 2-fold increase in their phosphorylation were considered *in vitro* phosphorylated. To perform clustering based on the phosphorylation pattern, the fold change for each site in a particular channel was scaled, i.e. 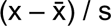, where s is the sample standard deviation and 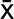 is the sample mean. Data were then plotted on a heat map and sites with missing values were eliminated. K-means clustering was then used to segregate sites with similar phosphorylation changes into clusters. Regular expressions were used to grep sites with certain amino acid sequences, such as those with or without a +1 Proline or those without a +1 Proline but with a +3 Lysine. To perform motif enrichment analysis, sequences of each cluster were inserted into the online tool WebLogo and plots generated were used in the results section presented here (Crooks et al., 2004). For motif enrichment analysis with IceLogo, the sequences in the cluster of interest were inserted as the positive set and sites in either the rest of the heat map or in the rest of the phospho-proteome were inserted as the background (Colaert et al., 2009). To match the CCR sites with the *in vitro* data, a column containing the gene name, the phosphorylated residue, and the location of that residue in the protein was added to the two tables. Rows with matching data in that column in both tables were considered *in vitro* targets of CDK1 with CCR endogenous phosphorylation.

## Supplementary information

